# Warming exacerbates fungal pathogenicity to drive physiological collapse and microbiome reorganization in octocorals

**DOI:** 10.64898/2026.08.12.744400

**Authors:** Matilde Marques, Francisca C. García, Yusuf C. El-Khaled, Pedro M. Cardoso, Erika P. Santoro, Neus Garcias-Bonet, Adam R. Barno, Matteo Monti, Gustavo A.S. Duarte, Helena D. M. Villela, Raquel S. Peixoto, Tina Keller-Costa, Rodrigo Costa

**Author notes:** These authors contributed equally to this work (Tina Keller-Costa, Rodrigo Costa).

## Abstract

Corals are increasingly threatened by climate change, yet the interplay between ocean warming and fungal infection remains experimentally underexplored. Here, we demonstrate that thermal stress exacerbates the pathogenicity of the fungus *Aspergillus sydowii*, driving physiological collapse and functional reorganization in the octocoral *Sclerophytum* sp. We deployed a 34-day mesocosm heat-stress experiment and quantified performance, mortality, and taxonomic and functional microbiome shifts. Combined heat and fungal exposure proved highly lethal, causing 45% mortality compared to 7% under heat exposure alone. Re-isolation from diseased nubbins implicates *A. sydowii* as the causal agent. Metagenomic profiling indicated that physiological collapse was underpinned by a functional transition from mutualism to antagonism in the microbiome. Specifically, combined heat and fungal stress triggered a depletion of ankyrin- and WD40-repeat proteins - hallmarks of symbiotic stability - concurrent with a surge in genes involved in fungal cell wall degradation and secondary metabolite biosynthesis. While surviving holobionts recovered full photosynthetic efficiency after combined stress, their microbiomes did not revert to baseline. Instead, they assembled into a taxonomically distinct configuration characterized by enrichments of sulfate-reducers (*Thermodesulfobacteriota*) and thermotolerant phototrophs (*Thermosynechococcales*). This suggests that despite the rapid photosystem recovery, the holobiont retained a complex legacy of thermal and biotic stress across both its internal chemical microenvironment and its microbiome. These findings highlight the decoupled recovery processes of different holobiont components, demonstrating that even though some corals may survive severe climate-driven disease, they emerge as ecologically reorganized entities.

## INTRODUCTION

Climate change is intensifying pathogenesis and increasing mortality across diverse natural and anthropogenic ecosystems, posing severe risks to both planetary and human health (*1–3*). In marine environments, the decline of corals and other foundation species threatens the ecological services upon which coastal ecosystems depend (*4*). As climate-sensitive marine ecosystems, coral reefs offer a valuable system for understanding complex disease dynamics in rapidly changing environments, particularly as climate change reshapes the composition of marine benthic communities worldwide (*5*, *6*).

Octocorals (class Octocorallia: comprising soft corals, blue corals, sea pens, and gorgonians/sea fans/sea whips) are both indicators and architects of shifting marine benthic ecosystems (*7*, *8*). While research has traditionally focused more often on tropical scleractinian corals of the class Hexacorallia, octocorals are increasingly recognized for their ecological importance as key habitat-forming organisms in a range of marine ecosystems. They occur across an exceptionally broad depth and thermal range, from shallow tropical reefs to the deepest and coldest waters across the globe (*9–11*). In recent years, reef ecosystems across the Caribbean and Western Atlantic, the Red Sea, and the tropical Indo-Pacific, have shifted from scleractinian coral dominance toward octocoral-dominated benthic assemblages, highlighting the resilience and recovery capacity of many tropical octocorals under environmental stress (*12–15*). Nevertheless, octocorals are also threatened by climate-driven disturbances (*16–18*). Marine heatwaves have caused widespread mortality in temperate regions where gorgonian forests play vital ecological roles (*19–21*). Thermal stress not only compromises holobiont physiology (including the breakdown of photosymbiosis where present) but also increases susceptibility to opportunistic and infectious diseases, further compromising octocoral health and survival (*16*, *22–24*).

The mechanistic basis of octocoral disease susceptibility remains poorly understood, particularly outside the gorgonian taxa historically associated with aspergillosis - a fungal disease affecting Gorgoniidae and Plexauridae families (*14*, *25*). This gap limits the ability to predict whether octocoral-dominated reefs will truly represent resilient future reef states or instead harbor emerging vulnerabilities to climate-driven disease. The filamentous fungus *Aspergillus sydowii* was initially identified as the probable etiological agent of aspergillosis in the gorgonian coral *Gorgonia ventalina* (*26*, *27*). However, its detection in both healthy and diseased octocorals was indicative of an opportunistic rather than strictly pathogenic role, modulated by host condition and environmental context (*28–30*). Other *Aspergillus* species have likewise been implicated in disease outbreaks exhibiting similar disease phenotypes in related gorgonian species (*28*, *29*).

As ocean temperatures continue to rise, the ecological balance of coral holobionts is increasingly disrupted, creating conditions that may favor opportunistic pathogens such as *A. sydowii*. This fungus exhibits a cosmopolitan distribution in marine environments (*31*, *32*), and both its growth and virulence are enhanced under elevated temperatures (*33*, *34*). Existing studies suggest that warming can alter host-pathogen interactions in octocorals by reducing the antifungal activity of coral crude extracts (*33*) and enhancing fungal protease activity (*35*). Additionally, infection with *A. sydowii* at ambient temperature has been shown to modulate host immune responses (*36*), while co-infection with copepods can trigger immune responses in octocorals (*37*). However, these studies have focused almost exclusively on *G. ventalina*, limiting our understanding of disease susceptibility across broader octocoral lineages. Moreover, how octocoral-associated microbiomes respond to interacting stressors remains unverified.

Here, we investigate the individual and combined effects of heat stress and fungal challenge on the tropical octocoral *Sclerophytum* sp., a soft coral that is becoming increasingly prevalent in Indo-Pacific benthic assemblages and is morphologically and taxonomically distinct from the gorgonian taxa commonly associated with aspergillosis. We hypothesize that fungal inoculation and elevated temperature each impair holobiont performance and restructure the microbiome, and that their combined effects will be additive or synergistic, facilitating fungal establishment and dysbiosis. By integrating phenotypic, physiological, and microbiome-level approaches, this study aims to disentangle how abiotic and biotic stressors interact to compromise octocoral health under contemporary ocean warming. By inoculating, detecting, and re-isolating *A. sydowii* from infected octocoral tissues, we provide experimental evidence consistent with a causal role for this opportunistic pathogen in compromising soft coral health. The pronounced decline observed under combined heat stress and fungal challenge demonstrates how synergistic interactions can destabilize coral holobionts and induce dysbiosis within their microbiomes, underscoring an emergent vulnerability of octocoral assemblages to climate-driven disease dynamics.

## MATERIALS AND METHODS

### Coral sampling, acclimation and fragmentation

Five specimens of the tropical octocoral *Sclerophytum* sp. were collected at Rose Reef, on the Saudi Arabian Red Sea coast (22°18’22.8”N 38°53’07.2”E). Samples were collected on November 8^th^, 2023, under a research permit with IBEC protocol number 22ibec003, using a hammer and chisel during SCUBA diving at depths ranging from 21.2 to 22.8 meters. The seawater temperature at the collection site was 30 °C. Corals were placed in plastic bags filled with surrounding seawater and immediately stored in cooler boxes also containing seawater. Within three hours of collection, the cooler boxes were transferred to laboratory facilities, where the coral specimens underwent initial acclimation to controlled conditions via gradual temperature adjustments through partial water exchanges before being placed in 300 L storage tanks with filtered seawater (industrial sand filters, 20 µm maximum filtration) maintained at 28 °C, as monitored by a Twin Temperature Controller (AquaMedic). The storage tank had a flow rate of 160 L/h, with water flow ensured by three pumps operating in random flow mode, regulated by EcoDrift controllers. A 12:12 h light:dark photoperiod was maintained using two Radion XR15 Pro lamps providing a maximum light intensity of 90-140 µmol photons m⁻² s⁻¹ (PAR). After 48 h of initial acclimation, each coral specimen was fragmented into 40 nubbins using sterile surgical scissors. Four nubbins from each specimen were immediately frozen in liquid nitrogen to serve as “baseline microbiome” references. The remaining nubbins were glued to sterile tiles and returned to the storage tank for a four-day healing period, allowing recovery from fragmentation stress. Following this healing phase, the coral nubbins were transferred to individual experimental tanks for a final seven-day experimental acclimation, marking the start of the experiment (designated as day 0).

### Mesocosm experimental setup and design

Four individual experimental tanks were assigned per treatment: control (C), fungus (F), temperature (T), and combined temperature + fungus (TF). These tanks were randomly placed within four 500 L water baths, in which uniform temperature was ensured by two pumps. Each experimental tank contained 8 L of microfiltered seawater (MFS), treated sequentially with sand filters (maximum 20 µm) and membrane filters (1 µm). Tanks were independently equipped with air pumps to ensure water flow and were completely individualized to prevent cross-contamination (Fig. 1A). Water temperature was controlled using a dual heating and cooling controller (D-DO Aquarium Solution). The system incorporated two heaters in each water bath and a sensor placed inside one of the experimental tanks. Partial water changes (20%) were conducted regularly every 1-3 days (Fig. 1B). Physical-chemical water parameters such as salinity, dissolved oxygen and pH were monitored every other day using a ProDSS 4-port digital sampling system (YSI). Salinity adjustments were made when required using deionized water.

**Fig. 1.**
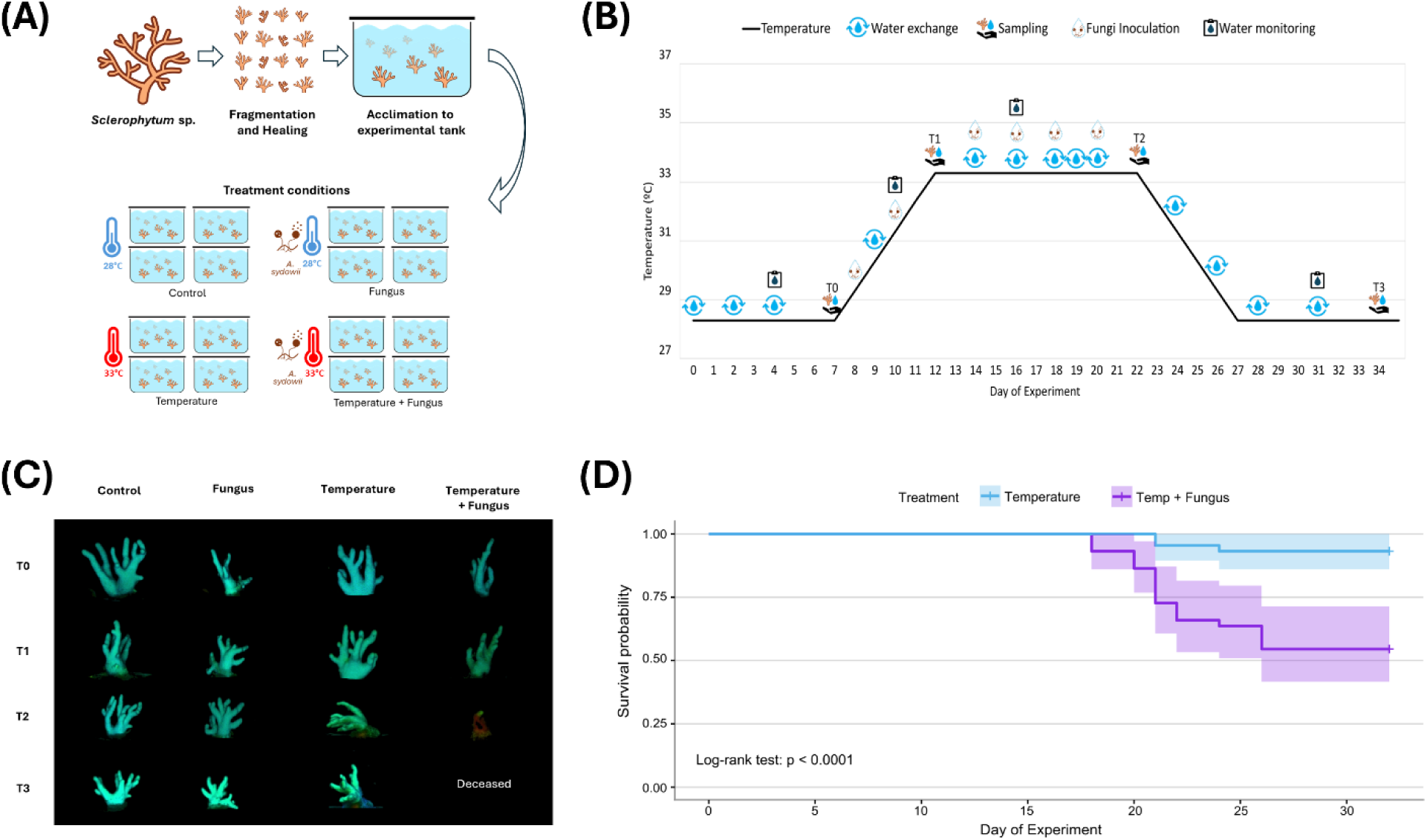
Experimental design and outcomes of heat and fungal stress experiment. (A) Workflow of sample processing from arrival at laboratory facilities to the start of the experiment, including experimental design. (B) Timeline of the experiment, detailing temperature regimes, timing of fungus inoculations, aquaria monitoring and maintenance, and sampling schedule. (C) Representative octocoral fragments from each treatment over time, summarizing the progression of stress responses. (D) Kaplan-Meier survival curves for octocorals exposed to Temperature vs Temperature + Fungus treatments. Shaded regions represent 95% confidence intervals.

The light cycle and intensity at the collection site and depth were monitored for 48 h with a Photosynthetically Active Radiation logger (Odyssey) deployed on site. To account for variability in turbidity, and the absence of an open circulation system, light intensity in the experimental tanks was reduced by approximately 50 PAR. This adjustment ensured a range of 81-100 PAR at peak intensity. Light was provided by four Radion XR15 Pro lamps per water bath and measured using a spherical sensor of the Universal Meter ULM-500 (Walz). A total of 173 *Sclerophytum* sp. nubbins were subjected to two thermal regimes: 28 °C as the control (C), and 33 °C to simulate temperature stress (T); and two inoculation conditions: fungus inoculation at 28 °C (F) and under temperature stress (33 °C; TF) (Fig. 1A). Two to three nubbins from each coral specimen were randomly placed in each experimental tank, resulting in 10-11 nubbins per tank.

Initially, all nubbins were maintained at 28 °C for seven days. For the treatments involving heat stress (T and TF), the temperature was increased by 1 °C per day until reaching 33 °C, which was maintained for 10 days. Subsequently, temperature was reduced back to 28 °C at the same rate, followed by a seven-day “recovery” period. Blank (see below) and fungus inoculations were administered every other day during both the temperature ramp-up and the 33 °C plateau period, and on equivalent days for the 28 °C treatments (C and F). For fungal treatments (F and TF), spores of *Aspergillus sydowii* were added to 400 mL of full-strength Marine Broth (Zobell) and incubated at 30 °C with continuous agitation at 100 rpm. After 10 to 15 days of incubation, cultures were filtered through 150 µm mesh to remove mycelium. The spore-containing filtrate was then centrifuged at 7,000 g for 15 min, and the resulting spore-containing pellet was resuspended in MFS - the same used in the experimental tanks. Spore concentrations were determined using a Neubauer hemocytometer and adjusted to 1.25×10^3^ spores per mL. The inoculant solution was applied directly to the water of each tank across all treatments, with water circulation ensuring even distribution. As a procedural control, an equivalent volume of MFS was used as a blank inoculant in treatments C and T. Thus, the four treatments used in this study were Control (C), fungus only (F), heat stress only (T), and combined heat stress and fungus (TF). Sampling was conducted simultaneously for all treatments at four time points: after acclimation (T0), following temperature increase and the initial phase of inoculations (T1), at the end of the stress plateau and/or inoculation period (T2), and after a recovery period (T3). A detailed schematic of the experimental design is presented in Fig. 1B. Holobiont health was routinely assessed through the photosynthetic efficiency of Symbiodiniaceae (see details below). At each sampling point, additional phenotypic, physiological and microbiome-related parameters indicative of holobiont health status were evaluated, along with water-associated microbiome composition (as detailed below).

### Monitoring of coral health and survival

Visual assessments for signs of bleaching, tissue loss, disease, or mortality were conducted daily. Coral phenotypic responses were evaluated using the Coral Health Chart as a reference (*38*) for sampling time point T2 (end of peak stress) in comparison with Control at T0, and the resulting data were processed as described in the Supplementary Material and Methods.

Dead nubbins were promptly preserved in liquid nitrogen for subsequent microbiome analyses (detailed below). Survival analysis was performed using the Kaplan-Meier method (*39*), with all nubbins that remained alive at the end of the experiment (day 32) treated as censored observations. Comparisons were restricted to the Temperature and Temperature + Fungus treatments, as these were the only groups exhibiting mortality during the experimental period. Kaplan-Meier survival curves were then fitted using the survival package (v3.8.3), and differences in survival distributions were assessed using the log-rank test (survdiff function).

### Pulse Amplitude Modulation (PAM) fluorometry as a proxy for holobiont health

The photosynthetic efficiency of the symbiont microalgae Symbiodiniaceae was used as a proxy for octocoral holobiont health. This method enabled continuous monitoring every two to three days using Pulse Amplitude Modulation (PAM) fluorometry. A submersible DIVING-PAM-II system (Walz GmbH, Effeltrich, Germany) equipped with a blue-emitting diode LED (peak emission at 474 nm) was used. To minimize non-photochemical quenching and avoid photoinhibition artifacts, samples were dark-adapted for at least one hour prior to measurement. The maximum quantum yield of photosystem II (*F_v_/F_m_*) was recorded using the following settings: measuring light intensity = 5; measuring light frequency = 3; gain = 2; damping = 2; and Electron-Transfer-Rate-Factor (ETR-Factor) = 1.

Temporal dynamics of *F_v_/F_m_* was monitored in up to four nubbins per tank, with replicates loss due to mortality. To evaluate physiological recovery trajectories without the confounding effects of mortality (analyzed separately using Kaplan-Meier survival curves), *F_v_/F_m_* values of zero (indicative of dead nubbins) were replaced with “NA” following the onset of mortality. Statistical model selection, linear mixed-effects modeling, and post hoc analyses are described in the Supplementary Material and Methods.

At each sampling time, fluorescence images were taken using an Imaging PAM (Walz GmbH, Effeltrich, Germany) following a dark adaptation period of 30 min. The Imaging PAM was equipped with an IMAG-K7 camera and an IMAG-MAX/L filter. Settings were standardized across all measurements as follows: aperture 4-2.8, measuring light intensity 6-7, gain 2, damping 2, and frequency 1. Data acquisition was performed with the ImagingWin V2.56zn software (Heinz Walz GmbH, Effeltrich, Germany).

### Physiological assessment: primary production by oxygen flux measurements

To assess oxygen fluxes, eight nubbins per treatment (two per tank) were tagged and used consistently throughout the experiment to quantify oxygen consumption in the dark (dark respiration, *R*_dark_) and oxygen production in the light (net photosynthesis, *P*_net_). Detailed incubation procedures, oxygen-flux calculations, and linear mixed-effects model analyses are provided in the Supplementary Material and Methods.

### Microbiome characterization via amplicon and metagenome sequencing

The structure and function of the octocoral-associated microbiome was assessed using a combination of 16S rRNA gene amplicon (taxonomic profiling of prokaryotic communities) and untargeted shotgun metagenome sequencing (microbiome functional profiling). Additionally, the molecular detection of *A. sydowii* in the samples was attempted using amplicon sequencing of the fungal internal transcribed spacer (ITS) region. For each treatment and sampling time, one nubbin per tank (from a distinct colony within the same treatment) was collected, yielding four biological replicates per treatment per time point. For each nubbin, a 0.5 g aliquot was aseptically removed using sterile pliers and a scalpel and immediately frozen in liquid nitrogen. Additionally, the “baseline microbiome” samples (see above) and 14 nubbins that died during the experiment were also included in microbiome assessment. Each nubbin was stored at −80 °C until total-community DNA (TC-DNA) extraction.

To assess the water-associated microbiome, approximately 1 L of seawater was collected from each tank at each sampling time (four replicates per treatment per time point) and stored overnight at 4 °C. Water samples were filtered through sterile 0.22 µm nitrocellulose membrane filters (MF-Millipore, 47 mm diameter) using a vacuum pump set to ≈15 cmHg. Filters were aseptically halved, and each portion was stored at −80 °C until TC-DNA extraction.

TC-DNA was extracted using the DNeasy PowerSoil Kit (Qiagen), following the manufacturer’s protocol with modifications to steps 3 and 4. These steps were replaced by three bead-beating cycles at 30 Hz, with one-minute intervals, using a TissueLyser II (Qiagen). Two negative controls were included and processed identically to the experimental samples: a “filter control” (extraction performed on a sterile nitrocellulose membrane filter) and a “kitome control” (extraction performed without biological material). The DNA concentration was quantified using a Qubit 2.0 Fluorometer High-Sensitivity DNA Kit (Fisher Scientific). DNA was stored at −20 °C until sent for sequencing.

Amplicon libraries targeting both bacterial and archaeal 16S rRNA genes and fungal internal transcribed spacer (ITS) regions were prepared following Earth Microbiome Project protocols. The hypervariable V4 region of the 16S rRNA gene was selectively amplified using primers 515F (5’–GTGYCAGCMGCCGCGGTAA–3’) and 806R (5’–GGACTACNVGGGTWTCTAAT–3’) (*40*, *41*), following the 16S Illumina Amplicon Protocol (https://earthmicrobiome.org/protocols-and-standards/16s/). For ITS region amplification, primers ITS1f (5’-CTTGGTCATTTAGAGGAAGTAA-3’) and ITS2 (5’-GCTGCGTTCTTCATCGATGC-3’) were used, following the methodology described in the ITS Illumina Amplicon Protocol (https://earthmicrobiome.org/protocols-and-standards/its/). Sequencing for both amplicons was conducted on an Illumina MiSeq platform at the Genomics Facility of the Gulbenkian Institute for Molecular Medicine (GIMM), Lisbon, Portugal, using paired-end reads: 2 × 300 bp for 16S rRNA gene and 2 × 250 bp for ITS. A total of 6,717,029 16S rRNA gene reads and 10,654,657 ITS reads were obtained, with per-sample read counts ranging from 2,353 to 175,077 and 452 to 319,599, respectively (Tables S1, S2). Two water samples (W-0TD and W-3TA) had only 2 reads in the 16S rRNA gene amplicon dataset and were excluded from analysis.

Shotgun metagenomic sequencing was performed to analyse the microbial communities of 24 samples collected at sampling time point T2 (corresponding to the end of the stress peak and/or inoculation period). These included four replicates per treatment, four “baseline microbiome” controls, and four samples from nubbins that died during the experiment under the combined temperature and fungal stress treatment (deceased temperature + fungus hereafter termed dTF). TC-DNA was sequenced at GIMM (Lisbon, Portugal). DNA libraries were prepared using the MGIEasy Fast FS DNA Library Prep Set (MGI, China) and sequenced using paired-end metagenome sequencing on an MGI G400 platform (large flow cell, PE150 cartridge). Raw sequencing data were demultiplexed by the sequencing facility prior to downstream analyses (Table S3).

### 16S rRNA gene amplicon data processing and quality control

Raw sequencing reads from 16S rRNA gene amplicon sequencing libraries were processed into amplicon sequence variants (ASVs) using the DADA2 (Divisive Amplicon Denoising Algorithm) R package (v1.30.0; (*42*)). Sequencing reads were trimmed at 10 bp and 120 bp (forward and reverse, respectively) at the 5’-end, and at 240 bp (forward) and 200 bp (reverse) at the 3’-end of each read. Subsequent filtering used default DADA2 parameters. Error rates were computed to identify unique sequences, and denoised reads were merged. Chimeric sequences were identified and removed, representing approximately 6.6% relative abundance of all reads. A total of 6,600 ASVs were initially inferred. Taxonomic assignment was performed using the naive Bayes algorithm (*43*) with the SILVA reference database v138.2 (*44*). Assignments were subsequently curated manually where necessary to comply with the List of Prokaryotic names with Standing in Nomenclature (LPSN) (*45*). ASVs not classified as prokaryotic at the domain level (9 unassigned and 34 assigned to Eukaryota), as well as those identified as chloroplasts (110 ASVs) or mitochondria (150 ASVs) were removed. Contaminant ASVs were identified via the two negative control samples: “filter control” (113 ASVs) and “kitome control” (28 ASVs). ASVs present in these controls were proportionally subtracted from the read counts of corresponding biological samples. Singleton ASVs (15 reads) were excluded to improve data robustness. ASVs lacking taxonomic classification from phylum to genus were queried against the NCBI nucleotide database using BLAST. This identified an additional 32 ASVs of mitochondrial origin, which were removed, totaling 724,057 reads. After all quality control and curation steps, 3,831,383 reads and 6,243 ASVs were retained across 144 samples. A detailed breakdown of read and ASV counts across each processing step is provided in Table S1.

### Microbial diversity and community composition analyses

Analyses of alpha and beta diversity, as well as taxonomic and differential composition of 16S rRNA gene data were performed in R (v4.4.3). Alpha diversity was assessed using ASV richness and the Shannon-Wiener diversity index following rarefaction to the minimum sequencing depth observed across samples (2,337 reads per sample; Fig. S1). Differences among treatments, sampling times, and biotopes were evaluated using non-parametric statistical tests. Beta diversity was assessed using Bray-Curtis dissimilarities computed from Hellinger-transformed, non-rarefied ASV counts (vegan package v2.6.10). Community composition was evaluated using canonical analysis of principal coordinates (CAP), PERMANOVA, and principal coordinates analysis (PCoA). PERMANOVA was performed with 9,999 permutations, with permutations constrained within experimental tanks to account for the non-independence arising from repeated sampling of the same tanks through time. Temporal changes in microbial community composition within *Sclerophytum* sp. were further assessed by analyzing pairwise PERMANOVA R² values across sampling time points.

Differentially abundant microbial taxa were identified using two complementary approaches, DESeq2 v1.46.0 (*46*) and MaAsLin3 (Multivariable Association with Linear Models) v0.99.16 (*47*). DESeq2 was used for pairwise differential abundance between treatments or time points, whereas MaAsLin3 was used to model Treatment, Sampling Time, and their interaction, while accounting for repeated measurements.

Detailed descriptions of data processing, statistical models, assumption testing, multiple-comparison corrections, model parameters, and software packages are provided in the Supplementary Material and Methods.

### Metagenome data processing and functional profiling

Unassembled reads were subjected to quality control and filtration using FastQC (v0.12.1; Kbase app v1.2.2) and Trimmomatic (v0.39; Kbase app v1.2.15) under default parameters within the KBase platform (*48*). To remove host and Symbiodiniaceae sequences, trimmed reads were mapped against a custom database using Bowtie2 (v2.4.1). The --un-conc-gz parameter was applied to retain paired-end reads that failed to align to the eukaryotic references. The mapping database comprised reference genomes from six non-gorgonian *Malacalcyonacea* hosts (GCA_982319615 *Sarcophyton elegans*, GCA_004324835 *Dendronephthya gigantea*, GCA_042919415 *Erythropodium caribaeorum*, GCA_025400075 *Phenganax stokvisi*, GCA_025400095 *P. marumi*, GCA_025434845 *P. subtilis*) alongside ten Symbiodiniaceae genomes. Free-living lineages, or those typically not associated with corals were excluded, while GCA_965643015 *Breviolum minutum*, GCA_977036015 *B. psygmophilum*, GCA_947184155 *Cladocopium goreaui*, GCA_963970005 *Durusdinium trenchii*, GCA_977109205 *Symbiodinium microadriaticum*, GCA_964212085 *S. pilosum*, GCA_965279495 *S. tridacnidarum*, GCA_009767595 *S. kawagutii*, GCA_905221605 *S. natans* and GCA_905231915 *S. necroappetens* were considered. Unmapped reads were assembled into metagenomic contigs using metaSPAdes (v3.15.3), with default parameters (Table S3). The resulting contigs were subjected to functional annotation based on clusters of orthologous groups of proteins (COGs) and protein families (Pfams) using our in-house Melange pipeline (https://sandragodinhosilva.github.io/melange/).

Diversity patterns in functional composition were visualized using PCoA based on Bray-Curtis dissimilarities calculated from the Hellinger-transformed count tables (vegan package v2.7.2). For each functional assessment, 90% confidence ellipses were drawn for each treatment level. A PERMANOVA with 999 permutations was used to test for overall differences among treatments. Pairwise PERMANOVA tests were performed for all treatment combinations, and significant comparisons were identified using *p* < 0.05.

Significant pairwise differences were further examined using Similarity Percentage (SIMPER) analysis (vegan package v2.7.2). For each significant pairwise comparison, the Hellinger-transformed count abundance profiles were subset to the two relevant groups, and SIMPER was used to identify the genes contributing most to between-group dissimilarity.

Secondary metabolite biosynthetic gene cluster (SM-BGC) compositions were analysed using abundance profiles generated with antiSMASH v8.0.4 (*49*) via the multiSMASH wrapper (10.5281/zenodo.8276143), using default parameters.

### Fungal re-isolation and DNA extraction

At the final sampling time point (T3), two nubbins from different colonies and tanks in the combined temperature and fungus (TF) treatment were stored at 4 °C in 0.9% NaCl (1:10 dilution, w/v). To attempt re-isolation of the fungus *A. sydowii*, each nubbin was macerated using a sterile mortar and pestle in 0.9% NaCl (1:10 w/v). The resulting macerate was transferred to a sterile tube containing 2 mm glass beads and vortexed for 1 min. Serial dilutions were further prepared in 0.9% NaCl, and 100 uL of each dilution was spread-plated onto Potato Dextrose Agar (PDA) supplemented with chloramphenicol and tetracycline (PDA + CT, 10 mg/L). Plates were incubated at 30 °C in the dark and monitored every three days for fungal colony growth. As a positive control, spores from the plate used to prepare the original inoculum for the experimental tanks were also plated.

Because all emerging fungal colonies presented consistent morphology, one representative colony from each coral nubbin and one colony from the positive control plate were selected for DNA-based identification. The selected colonies were cultured in Potato Dextrose Broth (PDB) at 30 °C with agitation at 110 rpm. After 12 days of incubation, the liquid cultures were centrifuged at 4,100 g for 15 min. Fungal pellets were flash-frozen in liquid nitrogen and macerated. For each sample, 0.25 g of macerate was used for DNA extraction using the DNeasy PowerSoil Kit (Qiagen), following the manufacturer’s protocol, except in step 3 in which 3 cycles of bead-beating using FastPrep program 6 for 30 sec, with 5 min ice baths intervals, were performed instead.

The ITS region was amplified for taxonomic identification using primers ITS1F (5′-CTTGGTCATTTAGAGGAAGTAA-3′; (*50*)) and ITS2R (5′-GCTGCGTTCTTCATCGATGC-3′; (*51*)). PCR reactions were prepared as earlier described (*52*). Thermal cycling consisted of an initial denaturation at 95 °C for 5 min, 35 cycles of 95 °C for 30 s, 55 °C for 30 s, 72 °C for 30 s and a final extension at 72 °C for 5 min. PCR products were purified using Sephadex G50 columns (GE Healthcare Bio-Science AB, Uppsala, Sweden), quantified with an Invitrogen Qubit 4 fluorometer (Fisher Scientific) using the Qubit dsDNA BR assay kit, and Sanger sequenced using the forward primer. Chromatograms were quality-checked and manually trimmed in BioEdit v7.0.5.3 (*53*).

### Phylogenetic analysis of Ascomycota ASVs and re-isolated fungi

ITS ASVs were obtained as described in the Supplementary Material and Methods, using DADA2 (v1.34.0) with taxonomic assignment performed against the UNITE reference database. This dataset was processed to enable targeted monitoring of *A. sydowii* across samples. Eight ASVs assigned to the *Ascomycota* phylum (Table S4) were selected for phylogenetic inference. The dataset included ten voucher ITS sequences from closely related *Aspergillus* species, the ITS sequence of the original *A. sydowii* strain used for inoculation, the above mentioned two ITS sequences from fungi reisolated from inoculated nubbins (TF treatment), and the ITS sequence of the positive control isolate, sequenced to confirm identity following multiple subcultures (confirmation isolates). Three *Basidiomycota* sequences were included as outgroup.

All sequences were aligned using MUSCLE (*54*), and the optimal evolutionary model was identified in MEGA12 (*55*) via the “Find Best DNA/Protein Models” function. The Kimura 2-parameter model with a discrete Gamma distribution (K2+G) was selected. A Maximum Likelihood phylogeny was inferred with 1,000 bootstrap replicates under partial deletion, removing sites with less than 85% coverage, resulting in an alignment of 25 sequences comprising 170 nucleotide positions. The final tree was visualized and annotated using iTOL (Interactive Tree Of Life) v7.1 (*56*) and refined in Inkscape (*57*).

## RESULTS

### Interactive effects of heat and fungal stress impair octocoral holobiont function

Nubbins exposed to elevated temperature generally exhibited visible signs of tissue deflation and stiffening (Fig. 1C). However, phenotypic assessments based on Coral Health Chart comparisons between T0 (acclimation) and T2 (end of peak stress) revealed no bleaching across treatments.

Even though hue shifts occurred, they showed no consistent treatment association. Mortality, characterized by tissue disintegration, occurred exclusively under temperature-related treatments (T and TF), reaching 7% under T and 45% TF (Fig. 1C, D). Kaplan-Meier survival analysis confirmed a significant difference in survival between these two treatments (*p* < 0.0001), with nubbins exposed to TF exhibiting markedly reduced survival relative to T alone.

The photosynthetic efficiency of Symbiodiniaceae (*F_v_/F_m_*), used as a proxy for holobiont health (*58*), exhibited distinct treatment-specific trajectories (Fig. 2A). A linear mixed-effects model revealed significant effects of Time (F = 86.87, *p* < 0.001), Treatment (F = 53.75, *p* < 0.001), and their interaction (F = 37.77, *p* < 0.001). C and F treatments maintained stable *F_v_/F_m_* values (≈0.65) throughout the experiment, with no significant temporal variation across T0 – T3 (Holm-adjusted pairwise comparisons, all *p*-adj. > 0.05; Fig. 2A), remaining within the range typically reported for healthy corals (*59*, *60*). In contrast, temperature increase (T) induced a pronounced decline at T2 relative to T0 and T1 (Holm-adjusted pairwise comparisons, all *p*-adj. < 0.001; Fig. 2A), coinciding with the onset of mortality near the end of the stress phase (Fig. 1D). When temperature increase was combined with fungus inoculations (TF treatment), the decline was steeper and mortality rose to 45%, beginning midway through the stress phase, with T2 values significantly lower than T0 and T1 (Holm-adjusted pairwise comparisons, all *p*-adj. < 0.001). At T3, *F_v_/F_m_* increased significantly relative to T2 in both T and TF treatments (Holm-adjusted pairwise comparisons, both *p*-adj. < 0.001). Under T, *F_v_/F_m_* at T3 did not differ from T0 or T1 (Holm-adjusted pairwise comparisons, *p*-adj. > 0.05). In contrast, under TF, *F_v_/F_m_* at T3 remained significantly lower than T0 (Holm-adjusted pairwise comparisons, *p*-adj. = 0.008). No significant differences in *F_v_/F_m_* among treatments were detected at T0 or T1 (Fig. 2B). However, by T2, differences were pronounced (Kruskal-Wallis test: χ² = 46.45, df = 3, *p* < 0.001; Fig. 2B).

**Fig. 2.**
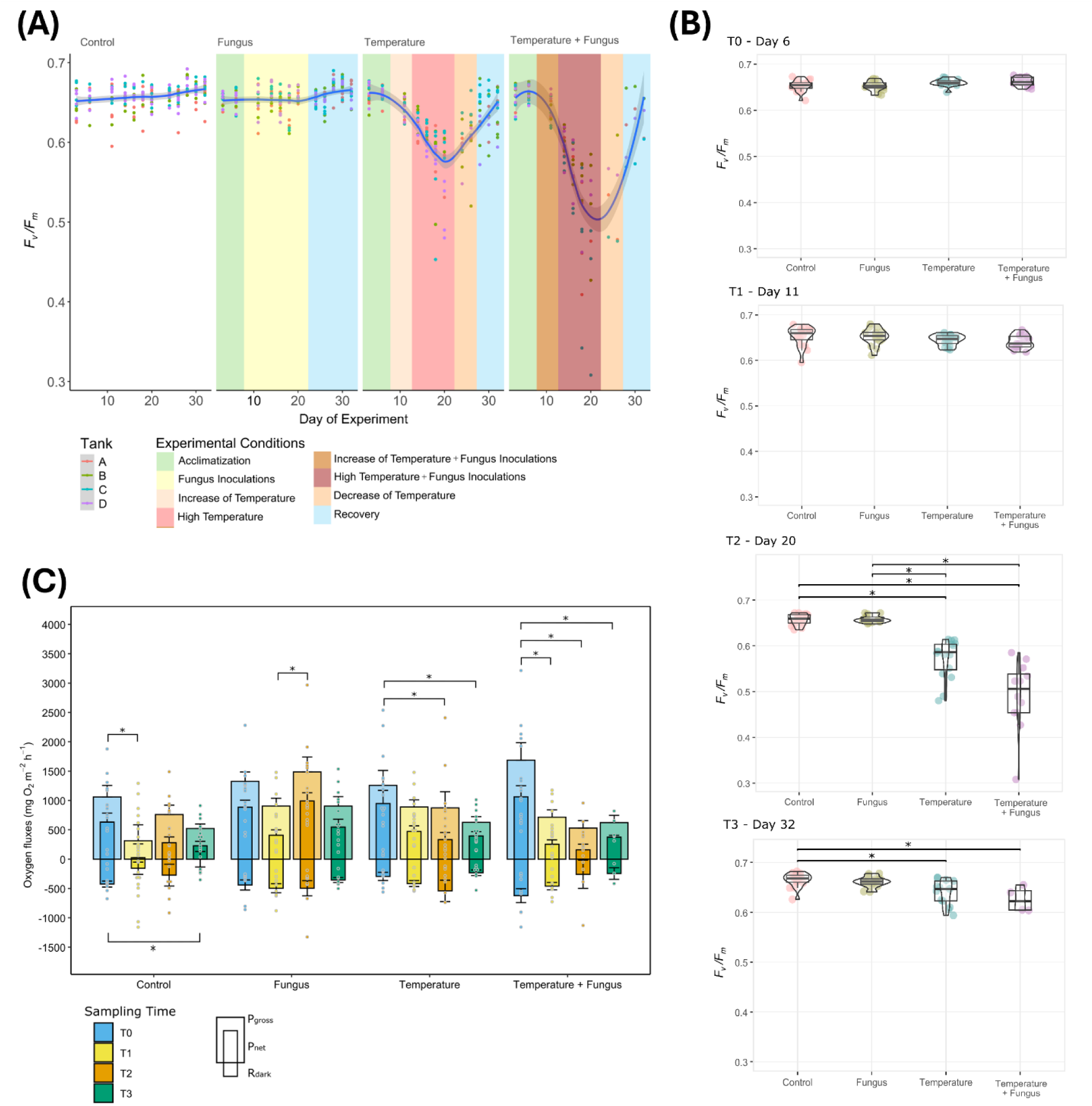
Physiological assessment of octocoral holobiont health. (A) Photosynthetic efficiency (*F_v_/F_m_*) of *Sclerophytum* sp. across treatments during heat and fungal stress experiment. Lines represent mean values and shaded bands show the 95% confidence interval round the fitted trend indicating the uncertainty of the estimate, with individual measurements shown as dots. (B) Statistical assessment of *F_v_/F_m_* at each sampling time: T0 – end of acclimation, T1 – beginning of stress period, T2 – peak stress, T3 – end of recovery phase. Significant differences between treatments (p < 0.05) were determined using Kruskal-Wallis tests followed by Dunn’s post hoc tests with Bonferroni correction, indicated by asterisks. (C) Gross photosynthesis (P_gross_), net photosynthesis (P_net_), and dark respiration (R_dark_) of *Sclerophytum* sp. over time for each treatment. Significant effects of time were assessed using type III ANOVA on linear mixed-effects models with Satterthwaite’s approximation, followed by Tukey-adjusted pairwise comparisons of estimated marginal means. Asterisks indicate significant differences (p < 0.05) between sampling times within each treatment for P_net_ (upper) and R_dark_ (lower). Pairwise comparisons for P_gross_ were identical to those observed for P_net_ under Control and Temperature + Fungus treatments. Error bars represent standard errors, and dots show individual measurements.

Bonferroni-corrected Dunn’s post-hoc tests confirmed significantly lower *F_v_/F_m_* values in T and TF compared with C and F (all *p*-adj. < 0.001). No differences were detected between C and F (*p*-adj. = 0.93) or between T and TF (*p*-adj. = 0.25). By T3, these differences diminished, with only C remaining significantly distinct from T and TF (Fig. 2B). Together, these results indicate that while thermal stress alone caused transient impairment of photosynthetic efficiency, the combined heat and fungal treatment resulted in a more severe functional impairment that was only partially reversible.

Independent measurements of oxygen flux mirrored PAM-based trends, confirming that functional impairment extended beyond photochemical efficiency to whole-holobiont metabolism (Fig. 2C). Stress treatments significantly influenced gross photosynthesis (*P*_gross_) and net photosynthesis (*P*_net_; *p*-adj. < 0.05), whereas dark respiration (*R*_dark_) was not significantly influenced. Within each treatment (Fig. 2C), pairwise comparisons indicated that both *P*_gross_ and *P*_net_ decreased between T0 and T1 in C (*p*-adj. < 0.05), and across multiple time points (T0 vs T1, T0 vsT2, T0 vs T3; all *p*-adj. < 0.05) in TF, reflecting a more dynamic response under combined stress. Additionally, *P*_net_ increased between T1 and T2 in F (*p*-adj. = 0.015), whereas in T a decrease occurred from T0 to T2 and from T0 to T3 (both *p*-adj. < 0.05). *R*_dark_ presented only a single significant difference which was under C, a decrease from T0 to T3 (*p*-adj. < 0.05). When grouped by time (Fig. S2), T2 was the only time point with significant differences among treatments. At this time point, *P*_net_ in F was significantly higher than in all other treatments, whereas T and TF did not differ from the control or from each other.

### Detection of *A. sydowii* reveals host colonization and persistence under stress

All fungal colonies re-isolated from experimental nubbins and control plates displayed similar morphology. Phylogenetic analysis of the ITS region confirmed that all isolates belonged to *A. sydowii*, showing 100% sequence similarity to the inoculated strain (Fig. 3A). In parallel, several unclassified Ascomycota ASVs detected via amplicon sequencing clustered closer to *A. sydowii* than to other *Aspergillus* species, forming a distinct clade we termed as the “*A. sydowii* clade”. Within this clade, ASVs 423 and 426 were consistently detected across multiple samples (Fig. 3B), with ASV 426 exhibiting the highest similarity to the inoculated strain (Fig. 3A). ASVs from the *A. sydowii* clade were predominantly associated with F and TF treatments, occurring in both seawater and *Sclerophytum* sp. microbiomes (Fig. 3B). Notably, ASV 426 and ASV 423 were also detected in deceased coral nubbins, further supporting their association with mortality following *A. sydowii* inoculation under heat stress.

**Fig. 3.**
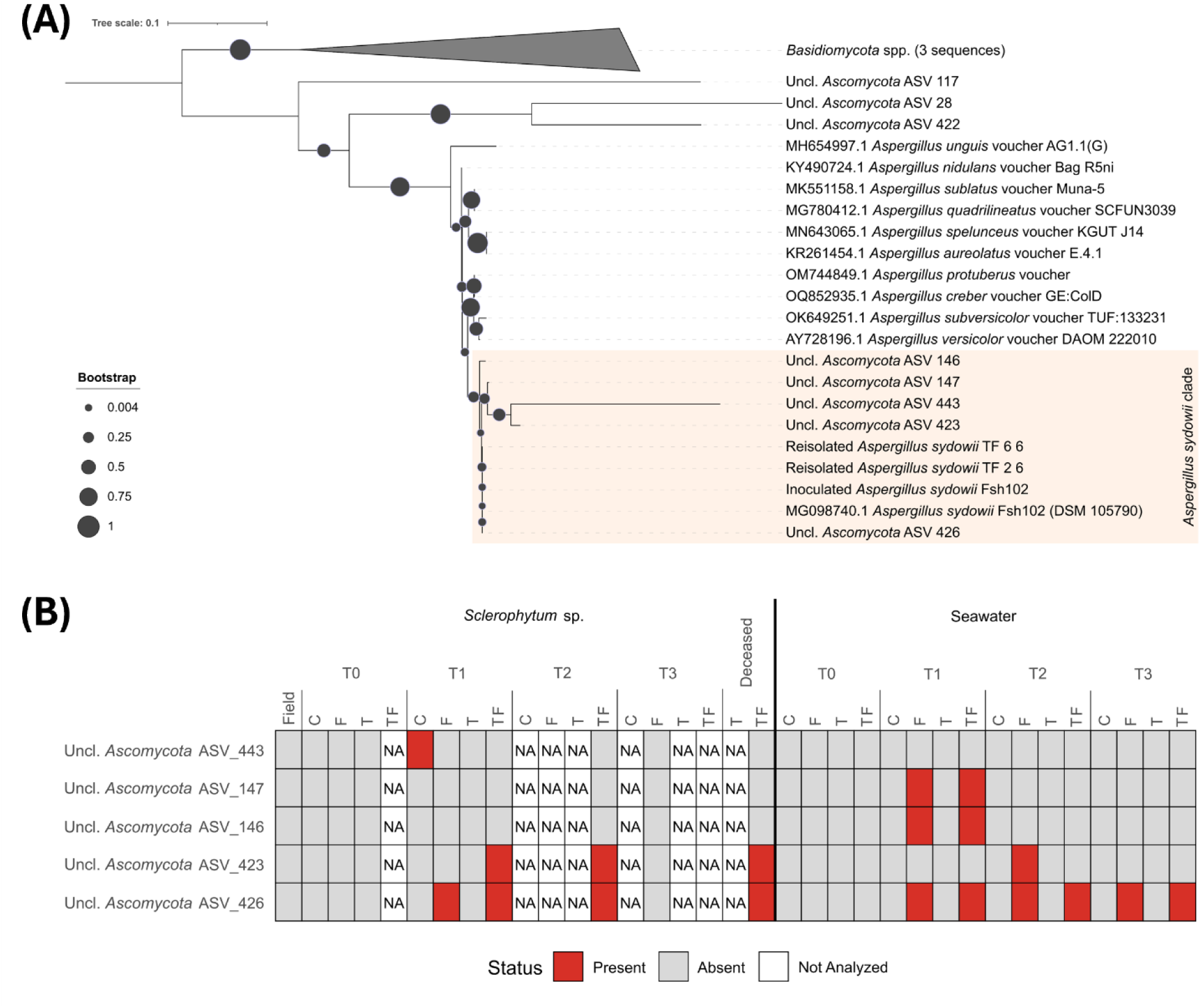
Molecular detection of *Aspergillus sydowii*. (A) Phylogenetic inference based on the ITS region of *Ascomycota* amplicon sequence variants (ASVs) and of re-isolated fungal strains. The tree includes eight *Ascomycota* ASVs, ten voucher sequences from closely related *Aspergillus* species, the sequences from the original *A. sydowii* strain purchased for the study (Fsh102, DSM 105790) and the corresponding inoculum used in the experiments (Inoculated *A. sydowii* Fsh102), and two sequences from re-isolated *A. sydowii* from deceased nubbins. Three Basidiomycota type strains were included as an outgroup. The tree was constructed using the Maximum-Likelihood algorithm with the Kimura 2-parameter model, with a discrete Gamma distribution (5 categories; +G, parameter = 1.7294). Partial deletion was used and all positions with less than 85% site coverage were eliminated, resulting in 170 positions in the final dataset. Filled circles indicate bootstrap support based on 1,000 replications. Branch lengths represent the number of substitutions per site (scale bar in the figure). ITS voucher accession numbers are shown before strain names. (B) Heatmap displaying presence/absence of *A. sydowii*-related ASVs across samples. “NA” indicates groups excluded from analysis due to insufficient ITS sequencing data in at least two samples per treatment after quality control and taxonomic filtering.

### Stress exposure drives shifts in prokaryotic community structure in octocorals

Alpha diversity measurements of ASV richness and diversity differed among biotopes (Fig. 4A). *Sclerophytum* sp. samples exhibited higher ASV richness and Shannon diversity than seawater (Wilcoxon test, *p* < 0.05 for both metrics; Table S5), with median richness of 244 ASVs in coral tissue compared with 77 ASVs in seawater. When testing for treatment- and time-dependent effects within each biotope, seawater exhibited significant variation in both ASV richness and Shannon diversity across Treatment × Sampling Time combinations (Kruskal-Wallis, *p* < 0.05). In contrast, no significant differences were detected within *Sclerophytum* sp. (Table S5).

**Fig. 4.**
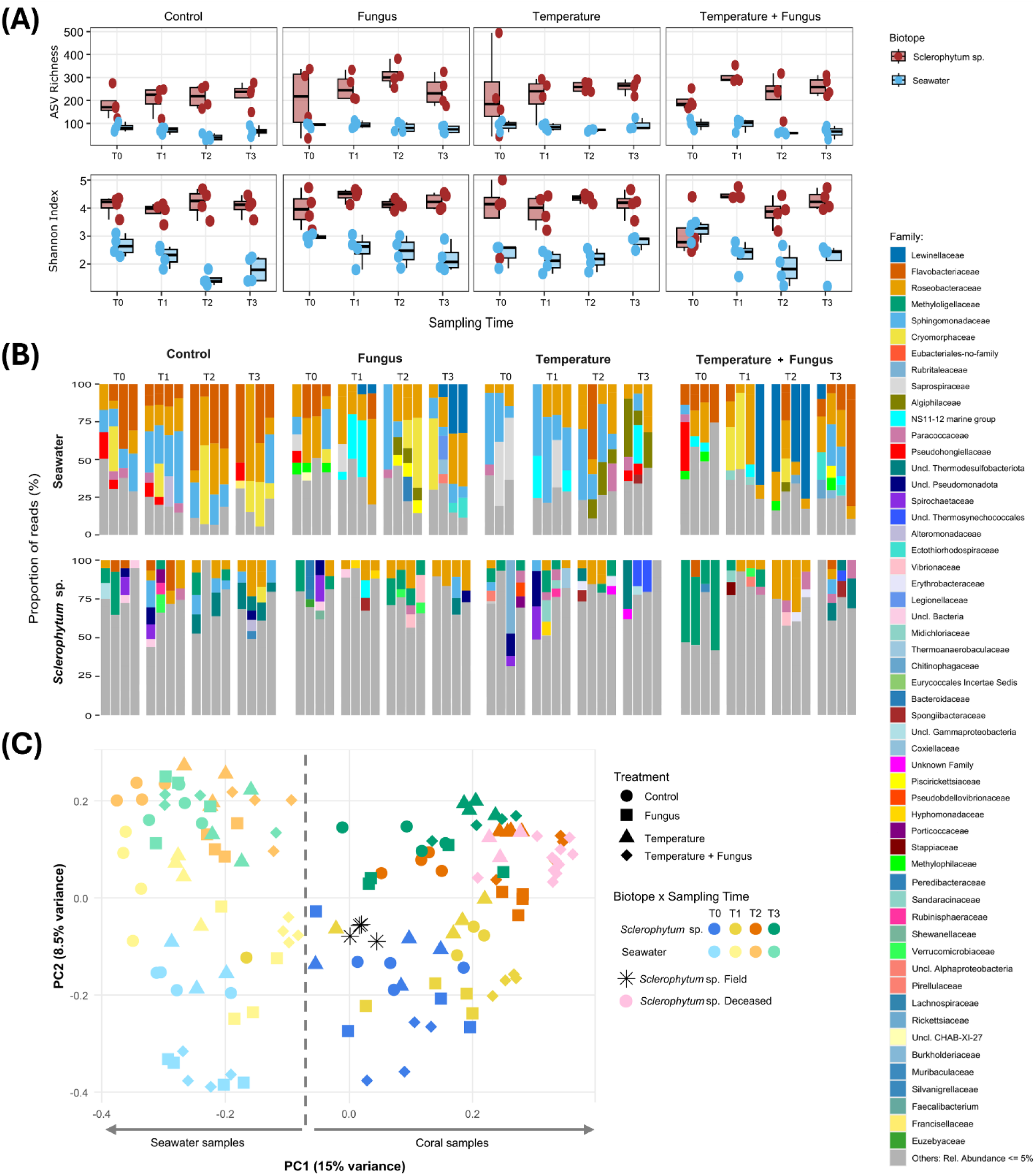
Prokaryotic community dynamics in *Sclerophytum* sp. and aquarium seawater throughout the experiment. (A) Alpha-diversity analysis of prokaryotic communities associated with *Sclerophytum* sp. (red) and seawater (blue). ASV richness and Shannon-Wiener diversity index are compared across treatments and sampling times, based on a rarefied dataset with a threshold of 2,337 sequences. Differences between biotopes were tested with Wilcoxon rank-sum tests, and differences within each biotope were evaluated using Kruskal-Wallis tests followed by Dunn’s post-hoc with Benjamini-Hochberg correction. Statistical summaries are provided in Table S5. (B) Family-level prokaryotic community composition of *Sclerophytum* sp. and seawater across time for each treatment. Relative abundances are presented for families contributing more than 5% of the total reads per sample. Taxa with lower relative abundances were grouped under “Other families”. (C) Multivariate analysis of prokaryotic community profiles. Principal coordinates analysis (PCoA) was performed on non-rarefied Hellinger-transformed abundance data, with treatments shown as shapes and sampling times as colours (dark = *Sclerophytum* sp., light = seawater). Canonical analysis of principal coordinates (CAP) was used to evaluate the independent effects, and a full-factorial PERMANOVA (999 permutations) tested the biotope × treatment × time interaction, followed by FDR-corrected pairwise tests. Statistical results are summarized in Table S5.

Consistent with the higher ASV richness and Shannon diversity observed in *Sclerophytum* sp., taxonomic profiles displayed a broader set of low-abundance families in corals than in seawater (Fig. 4B). Both biotopes exhibited pronounced compositional shifts over time and following stress exposure. These shifts were not accompanied by significant changes in alpha diversity, indicating compositional turnover without broad changes in richness. In *Sclerophytum* sp. (Fig. 4B), initial (T0) prokaryotic assemblages were dominated by *Methyloligellaceae* (synonym: *Hyphomicrobiaceae*, order *Hyphomicrobiales*), unclassified *Pseudomonadota* and *Spirochaetaceae*. Through time, stress treatments (F, T and TF) generally favored *Roseobacteraceae*, with this shift persisting after recovery in the T treatment. *Vibrionaceae* also emerged as dominant taxon at peak stress (T2) in both F and TF treatments, while *Paracoccaceae* became particularly associated with the TF treatment and persisted at T3. Notably, while the composition of prokaryotic communities under the F treatment loosely resembled that of the Control after recovery (T3), temperature-related treatments (T and TF) remained compositionally distinct, characterized by increased representation of unclassified *Thermodesulfobacteriota* and *Thermosynechococcales*. Shifts in seawater (Fig. 4B) are described in the Supplementary Results (Table S6).

Community ordination revealed clear separation of samples by biotope, with prokaryotic communities from seawater and *Sclerophytum* sp. forming distinct clusters across the first coordinate and distancing themselves from the baseline samples, which occupied a centred position in the ordination diagram (Fig. 4C). Indeed, biotope was found to explain the largest proportion of variation in beta diversity (*F₁* = 14.77, *p* = 0.001), with sampling time (*F₅* = 2.64, *p* = 0.001) and treatment (*F₄*= 2.27, *p* = 0.001) contributing additional effects according to Canonical Analysis of Principal Coordinates (Table S5). Coral samples at peak stress (T2) clustered close to recently deceased coral samples, indicating a stress-associated shift of the *Sclerophytum* microbiome toward communities typical of compromised coral health, particularly under treatments involving temperature increase (T and TF).

Across the full factorial design (Biotope × Treatment × Sampling Time), significant differences in prokaryotic community structure were found (PERMANOVA, *p* < 0.05), with 157 pairwise comparisons deemed significant, while 14 pairwise comparisons remained nonsignificant (Table S5). The TF treatment consistently produced the largest structural shifts from other groups, with divergence persisting into the recovery time point (T3).

Focusing on coral samples (Fig. 5A), stratified PERMANOVA revealed significant effects of Treatment (*R²* = 9.3%, *p* = 0.0001), Sampling Time (*R²* = 14.7%, *p* = 0.0001), and a significant Treatment × Sampling Time interaction (*R²* = 14.3%, *p* = 0.0337), demonstrating that stress responses were strongly time-dependent (Table S5). At T0, prior to stress exposure, microbial communities were largely similar across treatments, with most pairwise comparisons yielding non-significant results (*p*-adj. > 0.05). Divergence emerged immediately after initial stress exposure (T1) with all pairwise comparisons becoming significant (*p*-adj. < 0.05) and effect sizes increasing through T2, particularly for TF (*R²* = 0.42, *p*-adj. = 0.027). By T3, following the cessation of treatments, treatment-driven divergence remained evident, with significant pairwise differences (*p*-adj. < 0.05) and substantial effect sizes (*R²* = 0.27 - 0.38), particularly involving TF. Consistent with the ordination and pairwise PERMANOVA results, plotting the temporal trajectory of *R²* values showed that, although no quadratic terms were statistically significant, divergence among treatments peaked at T2, confirming this as the time of maximal microbiome dissimilarity (Fig. 5B). Temporal analyses within each treatment revealed substantial restructuring over time. While C exhibited moderate variability (*R²* ≈ 28%), stress-exposed treatments showed accelerated turnover (F ≈ 43%, T ≈ 40%, TF ≈ 53%), highlighting the cumulative impact of combined stressors (Table S5).

**Fig. 5.**
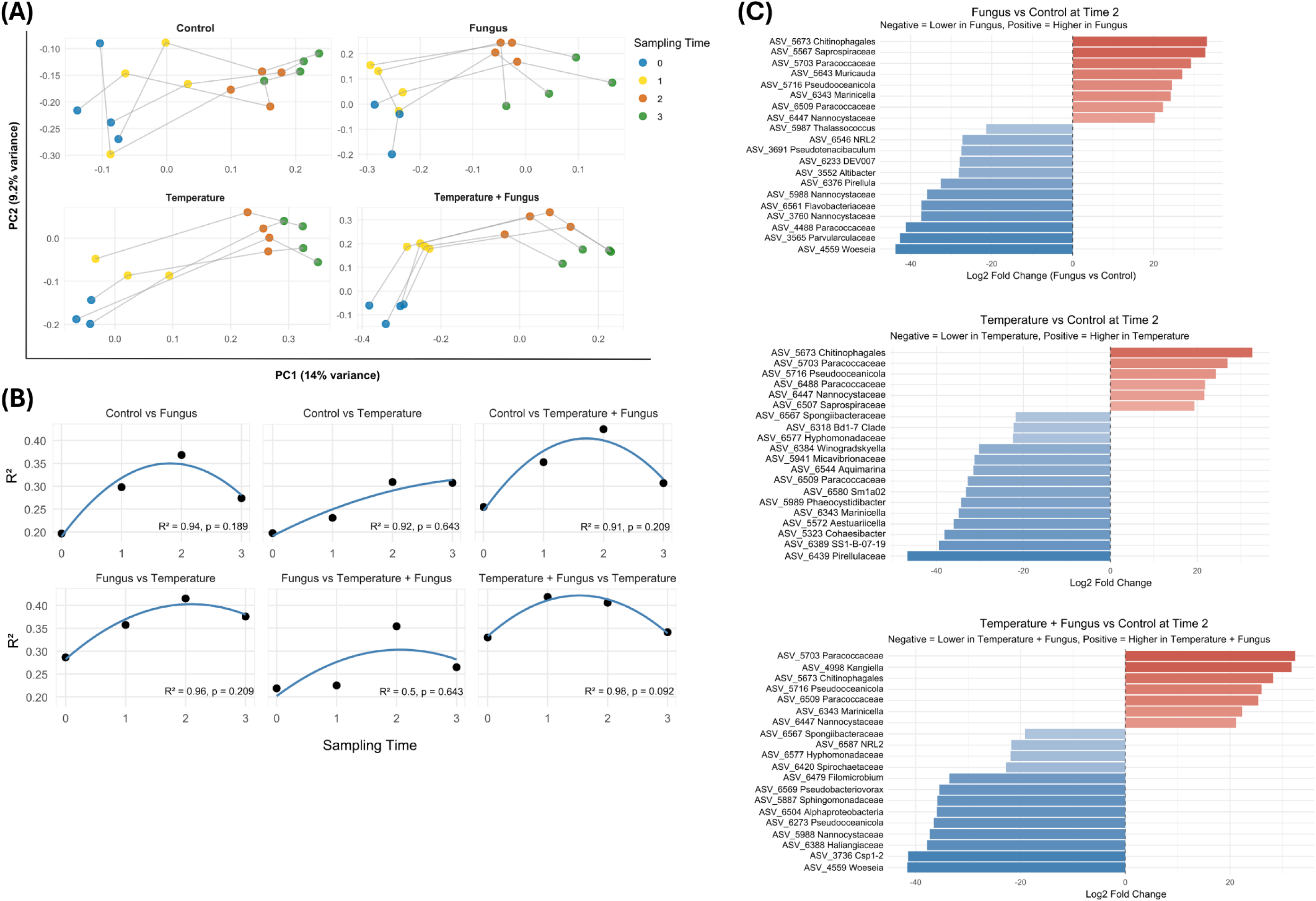
Within- and between-treatment prokaryotic community shifts in *Sclerophytum* sp. (A) Principal coordinates analysis (PCoA) of microbial community trajectories over time. Analysis was performed on non-rarefied Hellinger-transformed abundance data and plotted by treatment. Each line represents a replicate tank. (B) Temporal trends of pairwise PERMANOVA R² values for treatment comparisons. Points indicate observed R² values, and curves show quadratic regression fits. No quadratic terms were statistically significant (t-test on regression coefficients). (C) Top 20 differentially abundant ASVs at T2 for each stress treatment relative to the control, identified using DESeq2.

These results showed that while all treatments disturbed coral-associated communities, T and TF stress caused the greatest and most persistent divergence from the Control, with microbial communities failing to return to their original configuration during recovery.

### Stress-associated microbial responders in octocorals

Differential abundance analysis (DESeq2) at peak stress (T2) revealed clear prokaryotic signatures of stress exposure in the *Sclerophytum* holobiont (Fig. 5C; Tables S7-16), broadly consistent with the patterns observed in the taxonomic barplots (Fig. 4B). Several taxa responded consistently across stress conditions. Members of the order *Rhodobacterales*, particularly *Paracoccaceae* (e.g., ASVs 5703, 5707, 6509, 5714 and 6488) and *Pseudooceanicola* (ASV 5716), as well as *Chitinophagales* (ASV 5673), were consistently enriched across all stress treatments (F, T and TF). Conversely, multiple taxa were consistently reduced under T and TF, including *Spongiibacteraceae* (*Cellvibrionales*; ASV 6567) and *Hyphomonadaceae* (*Caulobacterales*; ASVs 6577, 6583 and 6579).

Besides these shared responses, several taxa exhibited treatment-specific patterns. At higher taxonomic levels (Tables S10-12, S15, S16), *Vibrionaceae* and *Vibrionales* were significantly enriched under both F and TF treatments, consistent with their dominance in the barplots at T2 (Fig. 4B). Moreover, T and TF treatments were characterized by an increase in *Phycisphaerales* at T2 that persisted in T3. Similarly, T was characterized by an increase in uncl. *Thermosynechococcales* at T2, which remained significantly enriched at T3. uncl. *Thermosynechococcales* also increased under TF at T3, alongside the decrease of *Hyphomonadaceae* and *Endozoicomonas* (Table S14).

Complementary MaAsLin3 modelling across all time points identified a set of ASVs significantly associated with treatment and time, partially overlapping with DESeq2 results, which are detailed in the Supplementary Results and Fig. S3.

### Stress-driven shift in microbial functions from symbiosis to degradation and defence

Principal Coordinates Analyses (PCoA) revealed consistent patterns of functional differentiation across both annotation frameworks (COGs and Pfams; Fig. 6A). In each case, samples from treatments involving elevated temperature (T, TF and dTF) clustered separately from the control and fungus treatments (C and F), indicating a strong and coherent temperature-driven restructuring of microbial functional composition. By contrast, baseline samples (prior to fragmentation and transfer to the experimental tanks) displayed variable placement in the ordination space and did not consistently align with any treatment cluster.

**Fig. 6.**
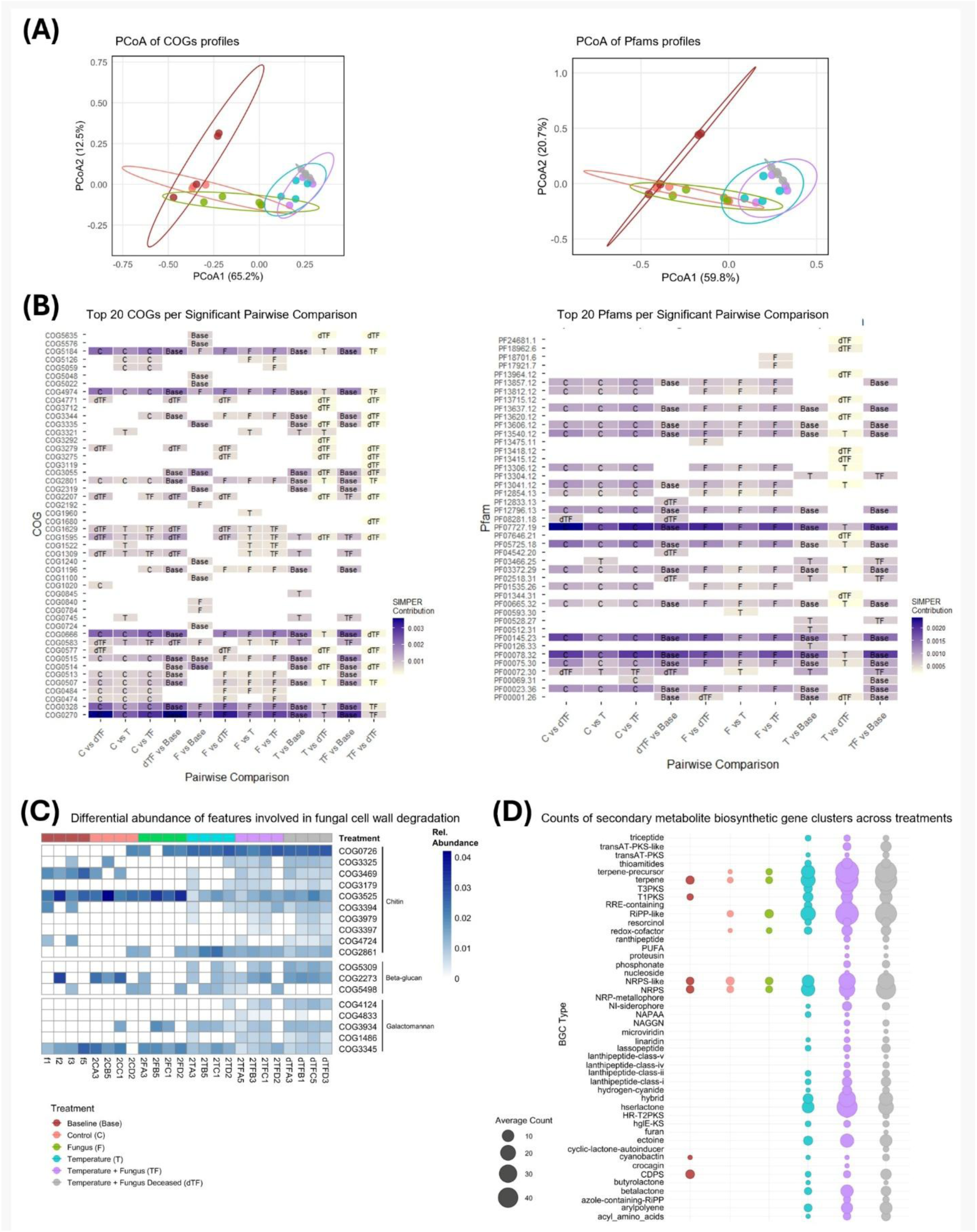
Functional profiling of the *Sclerophytum* microbiome at peak stress (T2). (A) Principal coordinates analysis (PCoA) of microbial functional composition based on Hellinger-transformed counts of COGs and Pfams. Ellipses represent 90% confidence intervals for each sample group. (B) SIMPER analysis of the top 20 contributing functions for significant pairwise comparisons. Tiles indicate the relative contribution of each function, with letters marking the group in which relative abundance was higher. Full functional descriptions for all COGs and Pfams are provided in Table S18. (C) Heatmap of chitin-related COGs across treatments. Complete descriptions of COG identifiers are listed in Table S18. (D) Biosynthetic gene cluster (BGC) profiles predicted for each sample group using antiSMASH.

PERMANOVA analyses (Table S17) confirmed significant overall differences in functional composition among treatment groups for both COG and Pfam annotations (all *p* = 0.001). Pairwise comparisons (Table S17) further revealed no significant differences between C and Baseline, C and F, or between T and TF, indicating that these treatment pairs exhibit similar functional profiles (all *p* > 0.05). In contrast, all other pairwise comparisons were significant (all *p* < 0.05). Differences between TF and dTF samples were significant only for COG annotations (*p* = 0.027).

To identify the functional features driving these differences, we performed SIMPER analyses for all significant pairwise contrasts (Fig. 6B). Across COG and Pfam datasets, microbial communities in C, F, and baseline samples were enriched in functions associated with stable host-microbe interactions and core cellular maintenance. These included protein-protein interaction domains such as ankyrin repeats (COG0666, PF13857, PF13637, PF13606, PF12796, PF00023), regulators of chromosome architecture (PF13540), as well as RNA and DNA processing functions (e.g., COG5184, COG0513, COG0328, PF07727). Notably, COG0270, which encodes a cytosine methylase involved in bacterial restriction-modification systems, was among the strongest contributors to these contrasts and was markedly depleted in T and TF.

In contrast, temperature-exposed treatments (T, TF, dTF) were enriched in functions linked to environmental sensing, transcriptional control, nutrient acquisition, and stress adaptation. These included the extreme heat-stress alternative sigma factor sigma24 (COG1595), broad-spectrum transcriptional regulators (COG1309, COG0583), and membrane-associated receptors and transport systems (COG1629, PF00528). The TF treatment was further enriched in additional transcriptional regulators, including the AraC (COG2207) and Lrp (COG1522) families.

Furthermore, the stress-adaptation response was most pronounced in dTF, which was uniquely enriched in virulence-associated systems (COG3275, COG3279, PF18962), specialized outer membrane receptors for iron scavenging (COG4771), and an antimicrobial peptide transport system (COG0577). This mortality-associated shift was further supported by the enrichment of secretion and invasion domains (PF18962), tissue-degrading carboxypeptidases (PF13715, PF13620), and galactose-binding domains (PF13418, PF13415), alongside a suite of response regulators and alternative sigma factors (PF08281, PF04542).

Focusing on COGs involved in the degradation of key fungal cell wall components such as chitin, β-glucans, and galactomannans, Hellinger-transformed abundance profiles (Fig. 6C) revealed a similar tiered response across treatments. Functions associated with degradation of the outer fungal cell wall, which is enriched in galactomannan and galactosaminogalactan, were particularly responsive to the combined stress (TF). Specifically, alpha- and beta-mannanases (COG4124, COG4833, COG3934, and COG1486) were selectively enriched in the combined treatments (TF and dTF), whereas COG3934 (endo-1,4-beta-mannosidase) was also enriched under T. Genes targeting the core fungal cell wall exhibited comparable patterns. Beta-glucan-degrading functions, including β-glucanases (COG5309 and COG2273), were enriched in TF, with COG2273 also showing elevated abundance in C. Chitin-related functions displayed a more differentiated response. COG3525, encoding beta-N-acetylhexosaminidases (EC 3.2.1.52), which can catalyse the hydrolysis of terminal non-reducing N-acetylglucosamine (GlcNAc) residues from chitin oligosaccharides (COS), was most abundant in baseline, C, and F, consistent with basal COS and aminoglycan turnover in stable coral microbiomes. In contrast, chitin-modifying functions such as COG0726 and COG2861 (polysaccharide deacetylases), and COG3469 (endochitinases, EC:3.2.1.14), presented higher abundance across heat-exposed treatments (T, TF, dTF), indicating that thermal stress (alone or combined) elevates microbial potential for chitin degradation via deacetylation and hydrolysis of internal glycosidic bonds. A subset of chitinolytic functions was overall restricted to the combined treatment (TF and dTF). These included COG3179 (GH19-type chitinase), COG3394 (COS deacetylase ChbG), COG3397 (CBM5/CBM33 carbohydrate-binding protein), and COG3979 (chitodextrinase), a suite of enzymes involved in the breakdown of chitin and its oligomeric intermediates.

AntiSMASH profiling of secondary metabolite BGCs further revealed heat- and fungus-driven shifts in secondary metabolite potential (Fig. 6D). Total BGC counts increased markedly in all heat-exposed treatments (T, TF, dTF) relative to C, F, and baseline. This expansion involved transAT-PKS-like, T1PKS, T3PKS, RiPP-like, thioamitide, homoserine lactone, betalactone, ectoine, arylpolyene, and acyl amino acid clusters, which are broadly associated with oxidative stress protection, membrane remodelling, antimicrobial activity, and stress signalling. A second set of BGCs appeared exclusively in TF and dTF, including PUFA, phosphonate, NAGGN, and multiple lanthipeptide-class clusters. These pathways, linked to antimicrobial peptides, membrane-active compounds, or interference molecules, suggest a shift toward opportunistic and antagonistic microbial strategies under combined thermal and fungal stress, particularly in fragments undergoing terminal decline.

## DISCUSSION

Octocorals are increasingly recognized as important organisms shaping marine ecosystems, contributing significantly to habitat complexity, biodiversity, and trophic networks. However, contrasting patterns have emerged regarding their sensitivity and adaptation to regional climate regimes. While many octocoral species have been identified as relatively resilient members of tropical coral reef ecosystems under climate change, often maintaining or increasing their abundance as scleractinian corals decline (*8*, *61*, *62*), growing evidence suggests that their responses are highly species- and context-dependent. Despite their ecological importance, studies investigating octocoral holobiont responses to interacting climate-related stressors remain scarce and are restricted to a limited number of species (*34*, *62–67*), leaving substantial gaps in our understanding of the mechanisms underpinning octocoral resilience and vulnerability under ongoing and future climate scenarios.

Whereas earlier work has largely focused on visual health metrics, culturable bacteria or host physiological and metabolic proxies, our study uniquely integrates multiple measures of holobiont performance with taxonomic and functional microbiome profiling. Our findings show that elevated temperature exacerbated the effects of fungal pathogen exposure in the soft coral *Sclerophytum* sp., resulting in persistent holobiont collapse without visible bleaching. The combined pathogen-temperature stress treatment resulted in disproportionate mortality, rapid loss of photosynthetic performance, and lasting taxonomic and functional microbiome reorganization, highlighting that octocoral vulnerability can remain cryptic and evade conventional reef health assessments (*16–18*, *68*).

### Physiological collapse under warming and disease

Thermal stress alone impaired photosynthetic efficiency and caused moderate mortality. When combined with fungal exposure, the stress caused a steeper decline in photosynthetic efficiency and quicker and more significant mortality. This synergy aligns with current evidence that climate change intensifies negative outcomes of host-pathogen interactions (*1*, *64*). Octocoral crude extracts have been shown to inhibit the growth of *A. sydowii*, but this capacity diminishes at high temperature (*33*). Moreover, metabolites produced by *A. sydowii* can impair the photophysiological performance of Symbiodiniaceae (*32*), and higher temperature regimes increase fungal protease activity (*35*), potentially weakening host immunity and accelerating tissue degradation. Together, these mechanisms provide a plausible basis for the heightened pathogenicity observed under combined thermal and fungal stress.

Consistent with the decline in photosynthetic efficiency, oxygen-flux measurements showed that temperature, particularly when combined with fungal exposure, reduced net photosynthesis, indicating impaired Symbiodiniaceae performance and diminished autotrophic energy acquisition. Although *A. sydowii* is widespread in marine environments (*31*, *32*), associations with octocorals have largely been restricted to gorgonian families such as *Gorgoniidae* and *Plexauridae*, which possess proteinaceous axial skeletons (*69*). Here, molecular detection and successful re-isolation of *A. sydowii* from the soft coral *Sclerophytum* sp. provide, to our knowledge, the first evidence of persistent colonization of a non-gorgonian octocoral host. The recurrent detection of *A. sydowii* in both living and perished tissues indicates that elevated temperature facilitates fungal establishment and persistence in association with the host. No classical signs of aspergillosis such as purple spots (*25*) were observed, suggesting context-dependent pathogenicity and expanding the recognized disease phenotype and spectrum.

### Stress legacies and the emergence of alternative microbiome states

Microbiome shifts were characterized primarily by taxonomic and functional reorganization rather than loss of microbial diversity. Microbial communities associated with *Sclerophytum* shifted gradually but followed consistent, stress-associated trajectories, with temperature and combined stress treatments inducing persistent deviations that failed to revert during the experimental “recovery” phase, even though holobiont photosynthetic performance returned to pre-disturbance levels. This pattern is compatible with the broader view that coral holobionts vary in their microbial response to environmental change, with some maintaining relatively stable microbiome configurations while others undergo more flexible microbial reassembly (*70*).

Such dynamics are consistent with the possibility of alternative microbial states, a well-established ecological concept across systems (*71–73*), in which communities reorganize into distinct configurations after disturbances. The greater divergence observed under combined stress at both taxonomic and functional levels supports the view that compounded stressors impose stronger selective pressures, increasing the likelihood of transitions toward novel microbial states. This indicates that the microbiome experienced a form of hysteresis, whereby recovery trajectories depend on historical conditions rather than current environmental states alone (*71*). As a result, the community failed to return to its original configuration. This interpretation is aligned with the recent proposals that coral holobionts can shift to “new baseline” microbial configurations following repeated or intense stress (*74*). Such baseline shifts could potentially have long-term implications, either through persistent alterations in microbial community composition or through stress-induced epigenetic modifications that influence future responses to environmental change (*75*, *76*). More broadly, these holobiont-level legacy effects may mirror ecosystem-scale patterns observed in Caribbean coral reefs, where historical disturbances constrain recovery trajectories and promote persistent alternative community states (*77*). Legacy effects of the disturbances induced in our experiment, particularly the compounded stress, were recognizable in the post-recovery prokaryotic communities, where taxa characteristic of thermotolerant (*78–80*) or sulfide-associated niches (*81*) increased in relative abundance, reflecting lasting alterations in the holobiont microhabitats.

During the stress phase, reduced photosynthetic efficiency suggests Symbiodiniaceae impairment, possibly opening ecological space for opportunistic colonizers such as thermotolerant cyanobacteria, including *Thermosynechococcales*, which subsequently increased in abundance by T3. Although the mechanisms underlying this shift cannot be resolved from our data, heat stress is known to destabilize nutrient exchange between corals and their algal symbionts and to shift holobiont metabolism toward alternative interactions among microbial partners (*82*, *83*).

Several bacterial taxa tightly associated with Symbiodiniaceae exhibited pronounced treatment- and time-dependent shifts. *Algiphilus*, *Phycisphaeraceae, Hyphomicrobium*, *Methylobacterium*, and *Sphingomonas*, among others, have been identified as bacteria associated to Symbiodiniaceae, with the latter three reported as intracellular Symbiodiniaceae symbionts (*84*). The overall decline of *Hyphomicrobiaceae* through time, combined with their detection among the most abundant taxa at T2 under control and fungal treatment, may therefore reflect progressive destabilization of bacterial functions that support coral-algal interactions under thermal and combined stress. These functions may include denitrification, C1 compound metabolism, and sulfur cycling, all of which play important roles in maintaining nutrient balance and metabolic homeostasis within the holobiont (*85–87*).

### Functional transition from symbiosis to degradation and defence

Thermal stress triggered a marked functional reorganization of the octocoral microbiome, extending well beyond shifts in taxonomic composition. Control and fungal treatments were enriched in functions associated with stable host-microbe interactions, intracellular signalling, and cellular homeostasis, features widely linked to symbiotic persistence in octocoral holobionts (*88*–*90*). In contrast, temperature-exposed treatments (alone and combined) showed enrichment of functions related to environmental sensing, transcriptional regulation, oxidative stress response, and nutrient acquisition, reflecting a shift toward a stress-responsive microbial state.

The most pronounced functional shift was observed under combined thermal and fungal stress, particularly in nubbins that ultimately died. These specimens were characterized by the enrichment of functions involved in fungal cell wall degradation, including pathways targeting galactomannans, β-glucans, and chitin. Notably, a suite of specialized chitinolytic functions, including ChbG, CBM5/CBM33, chitodextrinases and glycosyl hydrolase family 19 (GH19) endochitinases (*91*), was restricted to the combined-stress treatment. Together, these enzymes mediate successive steps in the breakdown of fungal cell wall polymers and their oligomeric intermediates, indicating an enhanced microbial capacity to exploit or antagonize structurally complex fungal substrates (*92*, *93*). The concurrent rise of chitinolytic taxa such as *Chitinophagales* and *Rhodothermales*, both recognized chitin degraders (*94*), strengthens this interpretation. Indeed, similar patterns have been described in plant rhizospheres, where chitinolytic bacteria and bioactive metabolites suppress fungal pathogens (*95*).

An alternative, and not mutually exclusive, explanation is that the enrichment of carbohydrate-degrading functions reflects the extensive host tissue deterioration observed under combined thermal and fungal stress. Corals in the combined treatment exhibited the greatest health decline and mortality, likely increasing the availability of organic substrates released from necrotic lesions, decaying tissue, and mucus-rich exudates (*96*). Under this scenario, the proliferation of carbohydrate-active enzymes and chitinolytic taxa may not solely represent a targeted response to fungal invasion, but also a broader shift toward opportunistic heterotrophy and organic matter recycling (*90*). Indeed, several of the enriched taxa, including members of *Chitinophagales*, are commonly associated with saprotrophic lifestyles and the degradation of complex biopolymers (*26*). The observed functional profile may therefore reflect the combined influence of fungal biomass turnover and increased access to host- and microbe-derived polysaccharides generated during tissue degradation.

This functional reorganization coincided with a pronounced depletion of symbiosis-associated features, including ankyrin- and WD40-repeat proteins, which are commonly implicated in host-microbe recognition, immune evasion, and symbiotic stability (*88*, *90*, *97–100*). Their decline, together with the loss of canonical coral-associated bacterial families such as *Spirochaetaceae* and *Hyphomicrobiaceae*, supports a breakdown of cooperative host-microbe interactions. In parallel, combined thermal and fungal stress was associated with a marked enrichment of BGCs, indicative of increased capacity for secondary metabolite production and intensified antagonism. Such shifts are frequently observed during microbiome dysbiosis, where the breakdown of stable host-associated interactions is accompanied by the proliferation of opportunistic taxa and an increasing importance of chemically mediated microbial interactions (*88*, *101*). Together, these changes, i.e., increased chitinases, expanded secondary metabolic potential and depletion of symbiosis hallmarks, indicate a microbiome shift toward a competitive- and defence-oriented functional regime. Stress-induced microbiome restructuring has been linked to broader ecosystem consequences, influencing interaction stability and functional persistence beyond the host organism itself (*102*).

This study demonstrates that the tropical octocoral *Sclerophytum* sp. responds to warming and disease through rapid physiological decline and sustained microbial reconfiguration. This contrasts with previous studies on tropical octocorals that have frequently inferred resilience based mainly on bleaching responses or short-term physiological recovery. Collectively, our results show that warming and fungal disease interact to drive holobiont collapse, resulting in an alternative microbiome state that may be functionally viable yet compositionally distinct. Such legacy effects may erode holobiont stability over repeated disturbances, reducing the capacity of octocorals to withstand the accelerating pace of climate driven disease events. Future research should quantify thermal and pathogenic effects across octocoral species and regions, test causality between specific microbial shifts and host decline, and evaluate whether microbial legacies alter population- and community-level dynamics. Additionally, it will be important to determine whether targeted microbiome-based interventions can reduce susceptibility to compounded stress. Recognizing and addressing these hidden vulnerabilities is essential for predicting and managing reef futures as climate change intensifies.

## Acknowledgments

We are grateful to the CMOR team for their invaluable technical and logistical support during the setup and maintenance of the aquarium systems. We also sincerely thank Miguel Viegas, Marco Casartelli, Ghaida Aljuaid, Phillipe Rosado, Giovanna Sabini-Leite, Corinne D’Anna, Bárbara Ribeiro, Axel Ortiz, Inês Raimundo and Laura Beenham for their assistance with laboratory work, monitoring, and sampling efforts.

## Author contributions

Conceptualization: M.Ma., Y.C.E.K., H.D.M.V., R.S.P., T.K.C., R.C.

Resources: Y.C.E.K., H.D.M.V., R.S.P., T.K.C., R.C.

Software: M.Ma., F.C.G.

Validation: M.Ma., F.C.G., Y.C.E.K.

Investigation: M.Ma., Y.C.E.K., P.M.C., E.P.S., N.G.B., A.B., M.Mo., G.A.S.D., H.D.M.V.

Formal analysis: M.Ma., F.C.G. Data curation: M.Ma., F.C.G. Visualization: M.Ma., F.C.G. Supervision: R.S.P., T.K.C., R.C.

Writing—original draft: M.Ma., R.C.

Writing—review & editing: M.Ma., F.C.G., Y.C.E.K., P.M.C., E.P.S., N.G.B., A.B., M.Mo., G.A.S.D., H.D.M.V., R.S.P., T.K.C., R.C.

Funding Acquisition: R.S.P., R.C.

Project administration: R.S.P., T.K.C., R.C.

## Data and materials availability

All data needed to evaluate the conclusions in the paper are present in the paper and/or the Supplementary Materials. The raw sequencing data of the 16S rRNA gene and ITS region amplicon datasets, provided as fastq files, have been deposited in NCBI’s Sequence Read Archive (SRA) under the BioProject accession number PRJNA1456944, and sample accession numbers SAMN57437115 - SAMN57437262. All metagenomes have been deposited in the European Nucleotide Archive (ENA) under the project accession number PRJEB111403 (ERP192018) and the sample accession numbers ERS29654212 - ERS29654235. The ITS region Sanger sequences of the fungal isolates have been deposited at NCBI GenBank under the accession numbers PZ374375 - PZ374377.

