## Supplemental Text and Figures for "Warming exacerbates fungal pathogenicity to drive physiological collapse and microbiome reorganization in octocorals"

Marques *et al.*

**This PDF file includes:**

Supplementary Text
Figs. S1 to S3
Legends for Tables S1 to S18
Citations to References 42, 46, 47, 103 to 107 (see Reference List in the Main File)

**Other Supplementary Material for this manuscript includes the following:**

Tables S1 to S18

**Supplementary Material and Methods**

Processing of coral visual monitoring data

Coral nubbins were photographed alongside the chart under standardized lighting conditions. Image processing was conducted in R using the imager package (v1.0.5). Regions of interest (ROIs) corresponding to coral tissue were manually delineated, while chart patch ROIs were predefined and applied to all images. Mean RGB values were extracted from each ROI and converted to CIELAB color space using convertColor function from the grDevices package (v4.4.3). To determine the closest color match, Euclidean distances in CIELAB space were calculated between each coral ROI and all chart patches. The chart patch with the minimum distance was assigned as the best-matching color. For each tank, a minimum of four nubbins were used to represent the overall phenotypic condition. Bleaching was defined as a reduction of at least two brightness units relative to the initial acclimation score (T0; e.g.: 6→4). Changes in hue category (B-E) without a corresponding drop in brightness were not classified as bleaching.

Statistical analysis of PAM fluorometry data

To evaluate overall trends, three linear models were initially compared using the lm() function in R (v4.4.3): a full interaction model (Time × Treatment), an additive model (Time + Treatment), and a time-only model. Model selection with ANOVA and Akaike's Information Criterion (AIC) identified the most complex model (interaction model) as the best fit. Incorporating this interaction structure, a linear mixed-effects model was then fitted (lme4 package, v1.1.37) for the data up until the end of the stress phase (T2; experimental days 0-22 of Fig. 1B), with Time, Treatment, and their interaction as fixed effects. Coral ID was included as a random intercept to account for repeated measurements of individual colonies. Estimated marginal means and pairwise comparisons were obtained using the emmeans package (v1.11.1). For within-treatment temporal contrasts among the four key experimental phases (day 6 - acclimation, day 11 - stress ramp up, day 20 - peak stress, and day 32 - recovery), Holm-adjusted $p$-values were applied to account for multiple comparisons. Treatment differences at each sampling time were evaluated separately. Following assessment of normality (Shapiro-Wilk test) and homogeneity of variances (Bartlett's test), Kruskal-Wallis tests were applied, followed by Dunn's post hoc tests with Bonferroni correction (FSA package, v0.9.6). All analysis were performed in RStudio (v2025.5.1.513; (*103*)).

Oxygen flux measurements and data analysis

Each nubbin was individually incubated in 160 mL airtight glass chambers (Weck) filled with its corresponding treatment water. To account for background planktonic metabolism, one additional chamber per tank was filled only with the respective water, run in parallel, and used as a blank control. All chambers were immersed in a temperature-controlled water bath, regulated by a thermostat-connected heater to match the respective treatment conditions. Each chamber was equipped with a magnetic stirrer (80 rpm), and stirring plates (Hanna Instruments) ensured constant mixing and uniform oxygen distribution.

Incubations consisted of a one-hour light incubation (c. 120 μmol photons $m^{-2}$ $s^{-1}$), immediately followed by a one-hour dark incubation. Dissolved oxygen concentrations were measured at the beginning and end of each incubation period using a SevenGo Duo handheld oxygen probe (Mettler Toledo). Net photosynthesis and respiration rates were calculated by subtracting initial from final oxygen concentrations, correcting for blank controls, and then normalizing to nubbin surface area (determined via the geometric method (*104*)), chamber volume, and incubation duration. The geometric method was selected because destructive alternatives (wax-dipping and dry-weight) were unsuitable given our need to retain samples for microbiome analyses, and buoyant-weight is equally confounded by tissue inflation/deflation. To reduce bias, surface-area measurements were carried out at consistent times of day under standardized conditions. Dark respiration is presented as a negative rate. Finally, gross photosynthesis ($P_{\text{gross}}$) rates were calculated as follows (equation I):

Equation I $$P\text{gross} = P\text{net} + |R\text{dark}|$$

To analyze the effects of treatment and time on each physiological parameter ($R_{\text{dark}}$, $P_{\text{net}}$ and $P_{\text{gross}}$), a linear mixed-effects modelling approach was employed (lme4 package, v1.1.37). The model included treatment, time and their interaction as fixed factors, and a random intercept for each coral nubbin to account for repeated measurements. Random-slope structures were evaluated but resulted in singular fits; therefore, the random-intercept model was retained. Model assumptions, including residual normality and homoscedasticity, were assessed using simulation-based residual diagnostics from the DHARMa package (v0.4.7). For each parameter, omnibus tests of fixed effects were performed with type III ANOVA using Satterthwaite's approximation for degrees of freedom, suitable for small sample sizes, via the lmerTest package (v3.1.3). P-values were adjusted using Benjamini-Hochberg correction to control the false discovery rate. When main effects or interactions were significant ($p$-adj. $< 0.05$), post-hoc pairwise comparisons of estimated marginal means were conducted using the emmeans package (v1.11.1) with Tukey adjustment, including (i) treatment contrasts within each sampling time point, and (ii) time contrasts within each treatment. Additionally, the overall treatment and overall time effects averaged across time points or treatments, were respectively assessed. All analyses were performed in R (v4.4.3, (*105*)).

<u>Alpha-diversity analysis</u>

For alpha-diversity analysis, sequencing depth was rarefied to 2,337 reads per sample to standardize coverage using the vegan package (v2.6.10). Alpha diversity was quantified using ASV richness and the Shannon-Wiener diversity index. Normality and homogeneity of variance were assessed with the Shapiro-Wilk and Levene's tests (car package v3.1.3), respectively. Shapiro-Wilk tests indicated that data deviated from normality ($p < 0.05$), while Levene's test showed that variances were generally equal across Treatment × Sampling Time combinations within each Biotope. Because the assumptions of parametric tests were not fully met across all groups, comparisons were conducted using non-parametric statistics. To test whether treatment and sampling time influenced alpha diversity within each biotope, a Kruskal-Wallis test using Treatment x Sampling Time as the grouping factor was performed. Significant results ($p$-value $<$ 0.05) were followed by Dunn's post-hoc test with Benjamini-Hochberg false-discovery rate correction (FSA package v0.9.6). Additionally, because only two biotopes were present, overall differences in alpha diversity between biotopes were tested using Wilcoxon rank-sum tests.

<u>Beta-diversity and community composition analysis</u>

Beta diversity was assessed using Bray-Curtis dissimilarities computed from Hellinger-transformed, non-rarefied ASV counts (vegan package v2.6.10). Homogeneity of multivariate dispersion was confirmed for Treatment, Sampling Time, and for each treatment across time using PERMDISP (9999 permutations), indicating no significant differences in within-group dispersion ($p < 0.05$). Canonical analysis of principal coordinates (CAP) was performed for each experimental factor (Biotope, Treatment, Sampling Time) to assess their independent contributions to community composition. Significance ($p < 0.05$) was assessed using 999 unrestricted permutations. In addition, an unrestricted, full-factorial PERMANOVA (Permutational Multivariate Analysis of Variance) with 999 permutations was performed using Bray-Curtis dissimilarities to evaluate the effect of the interaction term Biotope × Treatment × Sampling Time on community composition. Pairwise differences were further evaluated using pairwise PERMANOVA (pairwiseAdonis package v0.4.1), with *p*-values adjusted for multiple comparisons using the Benjamini-Hochberg false discovery rate correction. Community dissimilarities were visualized using principal coordinates analysis (PCoA) based on Bray-Curtis distances. For analyses restricted to *Sclerophytum* sp., samples were filtered to a minimum depth of 5,000 reads, and trajectories through time were plotted by treatment using tank ID as replicates. To evaluate temporal patterns in microbial community divergence, PERMANOVA $R^2$ values across pairwise treatment comparisons per sampling time were analyzed. Time was treated as a continuous variable, and both linear and quadratic regression models were fitted for each comparison to describe temporal trends. Quadratic significance was assessed using a t-test on the regression coefficient of the quadratic term ($Time^2$) in the model. Model outputs ($R^2$ values and *p*-values) were used for descriptive interpretation of temporal trends, including potential non-monotonic patterns.

Differential abundance analysis

To identify differentially abundant microbial taxa across treatments and time points, we employed two complementary statistical approaches: DESeq2 v1.46.0 (*46*) and MaAsLin3 (Multivariable Association with Linear Models) v0.99.16 (*47*). DESeq2 is optimized for count-based, pairwise comparisons and accounts for compositionality and sequencing depth, making it suitable for detecting strong differences in abundance between specific treatments or time points. In parallel, MaAsLin3 fits multivariable linear models with fixed effects for Treatment, Sampling Time, and their interaction, and includes Tank ID as a random effect. This framework accommodates repeated measures and continuous covariates, providing a complementary perspective on temporal and treatment-associated changes.

For DESeq2, low-abundance ASVs (total read counts ≤ 10 across all samples) were excluded to improve statistical power. The model incorporated fixed effects for Treatment, Sampling Time (discrete time points), and their interaction. Size factors were estimated using the poscounts (positive counts) method, and dispersions were fit using a parametric model. Differential abundance testing was performed using Wald tests, and p-values were adjusted for multiple testing using the Benjamini-Hochberg false discovery rate (FDR). Taxa with FDR < 0.05 were considered significantly differentially abundant. To interpret treatment effects at the end of peak stress and after recovery, contrasts were extracted to compare treatments within T2 and T3, respectively. The top 20 ASVs with the lowest adjusted p-values were visualized using DESeq2-normalized counts (mean ± standard error) across treatment groups. For MaAsLin3, microbial counts were normalized using total sum scaling (TSS) and log-transformed prior to modelling. Multivariable linear models were fitted with fixed effects for Treatment, Sampling Time, their interaction, and sequencing depth, while including Tank ID as a random effect to account for repeated measures. All variables were standardized to facilitate model convergence and interpretability. Statistical significance was assessed using FDR-adjusted p-values (Benjamini-Hochberg), with taxa meeting FDR < 0.1 considered differentially abundant across experimental conditions.

ITS amplicon data processing and taxonomic assignment

Raw ITS amplicon sequencing reads were processed into ASVs using DADA2 (v1.34.0; (*42*)), following a similar workflow as for 16S rRNA gene data. Primer sequences were removed with cutadapt (v4.0; (*106*)), and reads were quality-filtered using default DADA2 parameters. Error rates were estimated to infer unique sequences before merging paired reads, and chimeras were identified and removed, accounting for approximately 1% of total reads. In total, 552 ASVs were obtained.

Taxonomic assignment was performed against the UNITE general reference database for eukaryotes (v2, released on 19 Feb 2025; (*107*)). ASVs unclassified at Domain level, or assigned to *Metazoa* or *Viridiplantae* were filtered out, as were those detected in the kitome control. A detailed summary of read and ASV counts across each processing step is provided in Table S2.

**Supplementary Results**

Stress exposure as a driver of prokaryotic community structure in octocorals (continuation)

In seawater (Figure 4B; Supplementary Material File 1, Table S6), initial (T0) communities were dominated by *Flavobacteriaceae*, *Roseobacteraceae*, *Sphingomonadaceae*, and *Saprospiraceae*, with notable individual variability. Following initial stress exposure (T1), the *Lewinellaceae* family became dominant in both F and TF treatments. In TF, this increase continued from T1 to T2, reaching over 50% relative abundance in several samples at peak stress (T2). In contrast, F samples exhibited a temporary increase at T1, followed by a decline at T2 and a subsequent increase at T3 that exceeded T1 abundances. The *Algiphilaceae* family increased markedly at T2 across F, T and TF, particularly in F and T, and persisted as a dominant taxon only in the T treatment after recovery.

Taxonomic shifts and stress-associated microbial responders in octocorals (continuation)

Complementary MaAsLin3 modelling across all time points identified a set of ASVs with overall significant associations with treatment and time (Supplementary Material, Figure S2), with partial overlap with the DESeq2 results (Figure 5C). Early or peak-stress responders included *Nioella* (ASV 6407), which sharply increased at T1 in the F and TF treatments before declining, while *Porticoccus* (ASV 4596) peaked at T2 under both T and TF. *Cribrihabitans* (ASV 6416) also showed a T2 maximum abundance but only in the T treatment, with a subsequent decline at T3. A second group of ASVs exhibited cumulative increases under specific stress regimes. ASV 4599 (unclassified *Gammaproteobacteria*) increased continuously from T1 to T3 in the T treatment and also rose at T3 under F and TF. Likewise, ASV 6258 (*Hydrogenedensaceae*) increased consistently across all time points under T, and at later time points (T2-T3) under F and TF. Recovery-associated responses emerged also for a few taxa. ASV 3544 (Sm1a02, *Phycisphaerales*) increased only at T3 under T, while ASV 6515 (*Cyclobacteriaceae*) showed a delayed increase restricted to T3 under the combined TF treatment. ASV 3890 (UBA4486) displayed a contrasting pattern, increasing at T3 in F and TF but decreasing at the same time point under T. *Qipengyuania* (ASV 6465) instead consistently declined relative to the C, especially during stress exposure (T1 and T2) across all other treatments. Finally, *Marivita* (ASV 6563) increased steadily over time in the C and F treatments, suggesting sensitivity to thermal rather than fungal stress (Fig. S3).

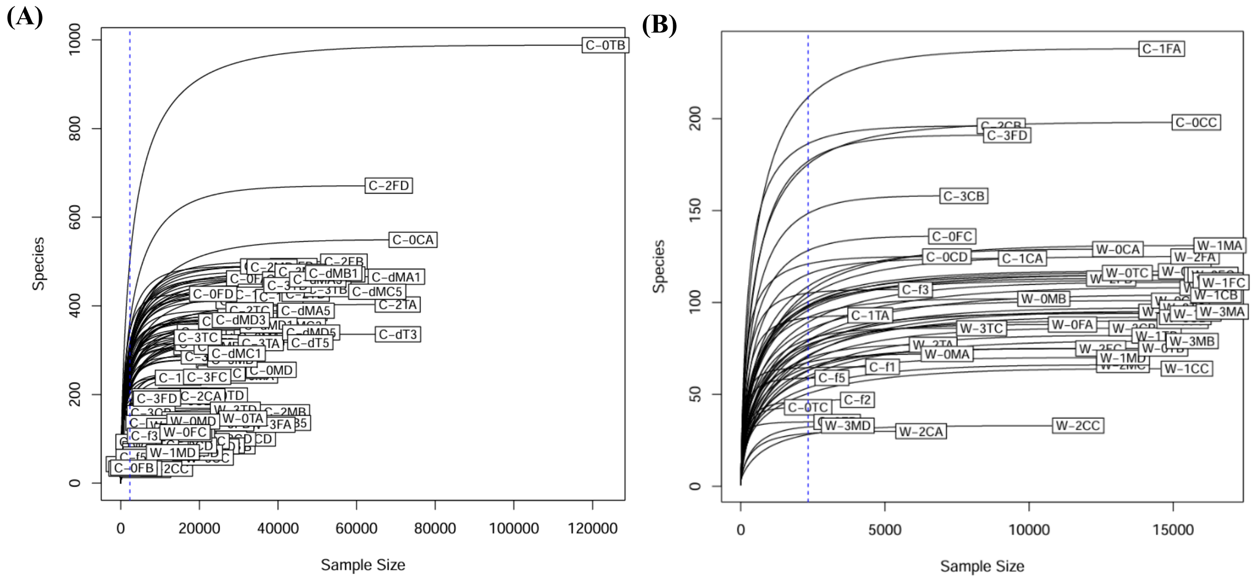


**Fig. S1.**

Rarefaction curves showing sequencing depth across samples. (A) Rarefaction curves for all 144 samples. (B) Rarefaction curves for the 50 samples with the lowest read counts. A blue horizontal line indicates the rarefaction threshold of 2,337 reads used for downstream alpha diversity analyses. Sample names correspond to those listed in Table S1.

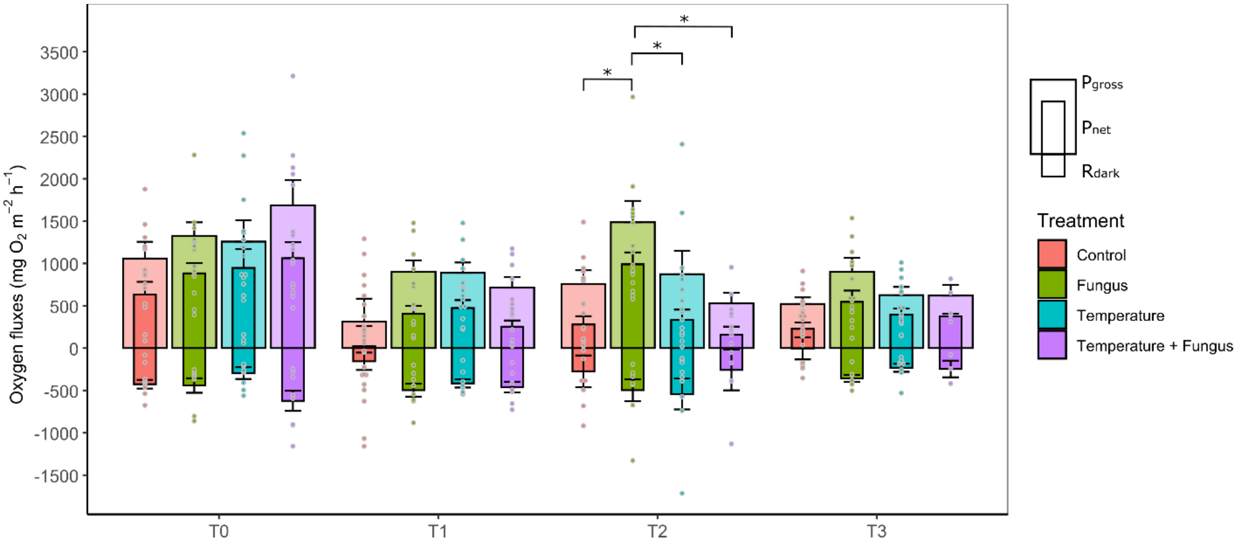


**Fig. S2.**

Gross photosynthesis ($P_{gross}$), net photosynthesis ($P_{net}$), and dark respiration ($R_{dark}$) of *Sclerophytum* sp. across treatments at each sampling time. Significant treatment effects were assessed using type III ANOVA on linear mixed-effects models with Satterthwaite's approximation, followed by Tukey-adjusted pairwise comparisons of estimated marginal means. Asterisks indicate significant differences ($p < 0.05$) between treatments within each sampling time for $P_{net}$ (upper) and $R_{dark}$ (lower). No significant pairwise differences were detected for $P_{gross}$. Error bars represent standard errors, and dots show individual measurements.

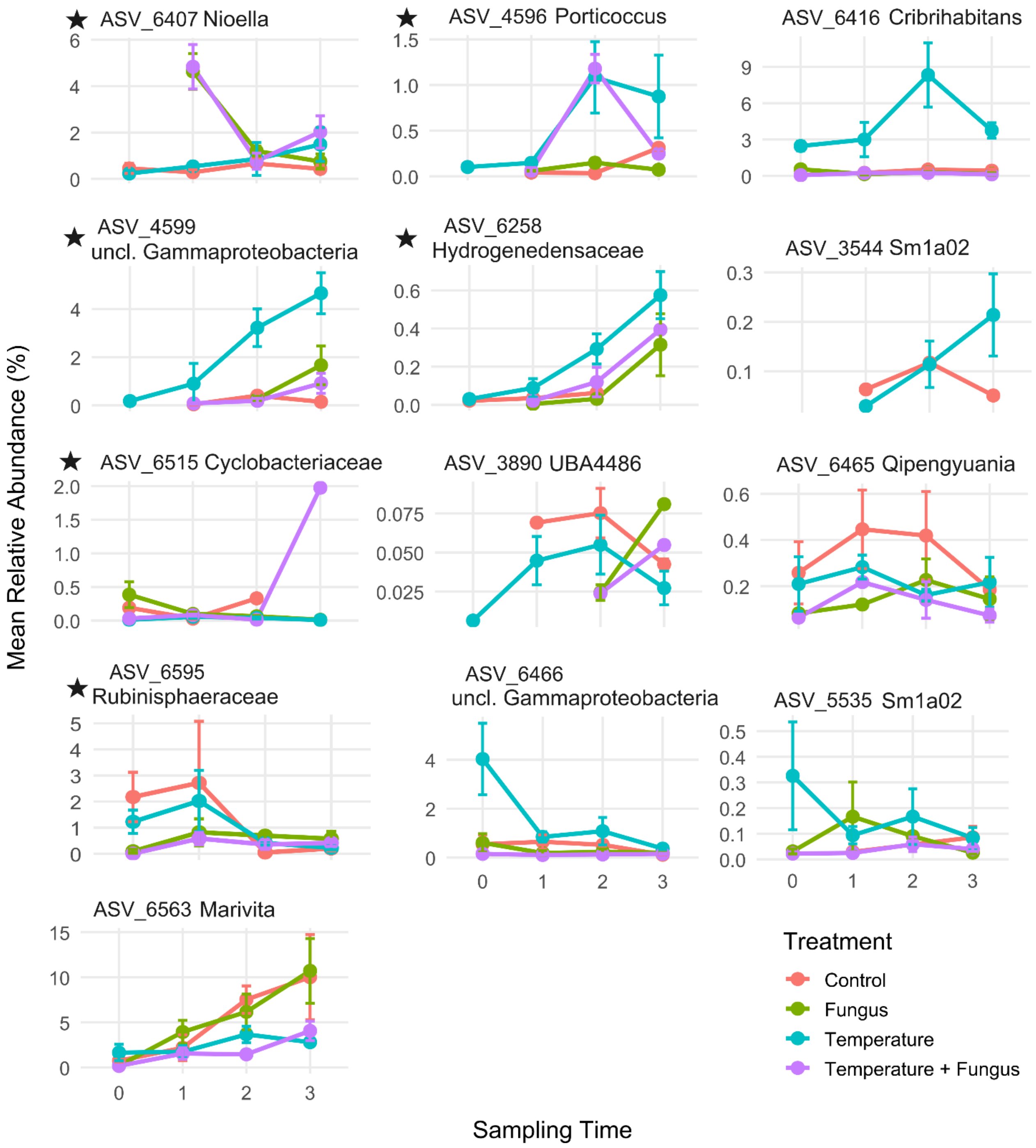


**Fig. S3.**
Temporal dynamics of significantly differentially abundant ASVs in *Sclerophytum* sp. Mean relative abundance (± standard error) of each ASV identified as significantly differentially abundant by MaAsLin3 is shown across all time points and treatments. ASVs that also overlap with the top 20 differentially abundant taxa identified by DESeq2 are indicated with an asterisk (*).

**Legends for Tables S1 to S18.**

**Table S1.**
Summary of the number of 16S rRNA gene reads and ASVs after each filtration/quality control step across the DADA2 pipeline.

**Table S2.**
Summary of the number of ITS reads and ASVs after each filtration/quality control step across the DADA2 pipeline.

**Table S3.**
General pre- and post-assembly features of the shotgun metagenomes sequenced at T2.

**Table S4.**
Final, quality filtered ASV table from the ITS amplicon sequencing data.

**Table S5.**
Statistical analysis of alpha- and beta-diversity metrics obtained for the 16S rRNA gene dataset.

**Table S6.**
Final, quality-filtered ASV table from the 16S rRNA gene amplicon sequencing data.

**Table S7.**
Differentially abundant ASVs identified by DESeq2 in *Sclerophytum* sp. at T2 - Fungus vs Control.

**Table S8.**
Differentially abundant ASVs identified by DESeq2 in *Sclerophytum* sp. at T2 - Temperature vs Control.

**Table S9.**
Differentially abundant ASVs identified by DESeq2 in *Sclerophytum* sp. at T2 - Temperature + Fungus vs Control.

**Table S10.**
Differentially abundant orders identified by DESeq2 in *Sclerophytum* sp. at T2 - Fungus vs Control.

**Table S11.**
Differentially abundant orders identified by DESeq2 in *Sclerophytum* sp. at T2 - Temperature vs Control.

**Table S11.**
Differentially abundant orders identified by DESeq2 in *Sclerophytum* sp. at T2 - Temperature vs Control.

**Table S12.**
Differentially abundant orders identified by DESeq2 in *Sclerophytum* sp. at T2 - Temperature + Fungus vs Control.

**Table S13.**
Differentially abundant ASVs identified by DESeq2 in *Sclerophytum* sp. at T3 - Temperature vs Control.

**Table S14.**
Differentially abundant ASVs identified by DESeq2 in *Sclerophytum* sp. at T3 - Temperature + Fungus vs Control.

**Table S15.**
Differentially abundant orders identified by DESeq2 in *Sclerophytum* sp. at T3 - Temperature vs Control.

**Table S16.**
Differentially abundant orders identified by DESeq2 in *Sclerophytum* sp. at T3 - Temperature + Fungus vs Control.

**Table S17.**
One-Way PERMANOVA of COG and PFAM functional profiles across treatments at peak stress (T2).

**Table S18.**
Description of the individual COG and PFAM entries displayed in Figures 6 B and C.
